# Lysine Crotonylation Acts as an Epigenetic Switch for Glutamate Neurotransmission and Spatial Memory

**DOI:** 10.1101/2025.02.19.639114

**Authors:** Utsav Mukherjee, Budhaditya Basu, Pratyush Suryavanshi, Stacy E. Beyer, Saaman Ghodsi, Nathan Robillard, Yann Vanrobaeys, Eric B. Taylor, Joseph Glykys, Ted Abel, Snehajyoti Chatterjee

## Abstract

Histone post-translational modifications (PTMs), particularly lysine acetylation (Kac), are critical epigenetic regulators of transcriptional programs underlying long-term memory consolidation. Beyond Kac, several other non-acetyl acylations have been identified with the ability to regulate transcription; however, their role in memory consolidation remains unknown. Here, we identify histone lysine crotonylation (Kcr) as a molecular switch for long-term memory and glutamatergic neurotransmission. We find that spatial learning induces distinct spatiotemporal patterns of Kcr across hippocampal subregions, and that Kcr stimulates learning-induced gene expression. Through genetic and pharmacological manipulations, we show that reducing hippocampal Kcr levels impairs memory, while increasing Kcr enhances long-term memory. Single-nuclei transcriptome and chromatin accessibility analyses reveals that Kcr facilitates activation of genes regulating glutamate signaling. Cell-cell communication analysis further infers that Kcr enhancement strengthens glutamatergic signaling within principal hippocampal neurons. Real-time fluorescence imaging with genetically encoded sensors functionally validates our multiomics and computational findings—demonstrating the role of Kcr in regulating presynaptic glutamate release and neuronal activity. In summary, our findings elucidate a novel mechanism underlying long-term memory consolidation and excitatory neurotransmission, linking epigenetic events to synaptic function and behavior.

## Introduction

Histone PTMs play a key role in the epigenetic regulation of gene expression during memory consolidation^1-5^. Short-chain acyl modifications on histone lysine residues are well-known PTMs that regulate several biological processes^6-8^. Among these, histone lysine acetylation (Kac) has been extensively studied for its role in memory storage. Altering histone Kac levels by selectively manipulating the ‘writers’ (lysine acetyltransferases, KATs) and ‘erasers’ (lysine deacetylases, KDACs) of histone Kac has been shown to impact hippocampal memory storage^2,9,10^. Interestingly, studies have revealed the promiscuous nature of the majority of the writers and erasers for histone Kac— demonstrating that these enzymes can also catalyze the incorporation or removal of non-acetyl acyl moieties on histone lysine residues^6,11,12^. This raises the question of whether the impact of manipulating KATs and KDACs on memory consolidation, as demonstrated in previous studies^13-18^, could be attributed to underlying changes in non-acetyl histone acylations besides Kac. Elucidating the functional relevance of non-acetyl histone acylations in long-term memory is therefore essential for advancing our conceptual understanding of epigenetic mechanisms underlying memory consolidation.

Advancements in mass spectrometry and biochemical approaches have led to the identification of several novel histone lysine acylations that are structurally distinct from histone Kac^19,20^. Among these, histone lysine crotonylation (Kcr), an evolutionarily conserved non-acetyl acylation, has gained significant attention in recent years for its role in transcription regulation^20-26^. Kcr is enriched on active gene promoters^20^, and Kcr-mediated regulation of gene expression has been implicated in diverse biological functions, including DNA damage repair^21^, acute kidney injury^25^, spermatogenesis^22^, nerve-injury-induced-neuropathic pain^26^, and neural stem cell differentiation^23^. Interestingly, studies in cell-free systems have demonstrated that the lysine acyltransferase p300 stimulates histone Kcr-mediated transcription to a significantly higher magnitude than histone Kac^12^. Furthermore, several histone ‘reader’ proteins—initially characterized as Kac readers— exhibit a higher binding affinity for crotonylated lysine residues compared to acetylated lysine moieties, and this preferential binding is thought to underlie the ability of the readers to regulate transcription^27^. We and others have shown that precisely timed transcriptional events, induced upon spatial learning, are critical for long-term memory consolidation^28-31^. Although Kcr has been well established as a transcriptional activator, its physiological relevance to memory consolidation—and signatures of Kcr-dependent gene expression programs that underlie hippocampal memory storage—remain elusive.

The availability of metabolic precursors dictates the landscape of histone acylations^6^. Accordingly, changes in intracellular crotonyl-CoA levels have been shown to alter histone Kcr levels, with direct consequences in gene transcription. Increasing crotonyl-CoA levels through crotonate administration elevates Kcr and stimulates gene transcription^12,23^, whereas depletion of Kcr via manipulation of the crotonyl-CoA hydratase chromodomain Y-like protein (CDYL) represses transcription^24^. Interestingly, in addition to writers and erasers, histone Kac and histone Kcr share enzymes that synthesize their respective metabolic precursors. The enzyme Acetyl-CoA synthetase 2 (ACSS2) synthesizes both acetyl CoA and crotonyl-CoA^12,32^. We previously demonstrated that ACSS2 is a critical regulator of neuronal histone Kac and spatial memory consolidation in the hippocampus^33^. Silencing ACSS2 levels reduces histone Kac and impairs activity-dependent transcription of memory-related genes^33^. Notably, knockdown of ACSS2 has also been shown to reduce Kcr levels and repress Kcr-regulated inflammatory gene transcription^12^. However, whether ACSS2-mediated regulation of histone Kcr constitutes a critical molecular mechanism underlying long-term memory consolidation remains unknown. Altogether, these findings raise the possibility that metabolic coupling of crotonyl-CoA availability to Kcr-dependent chromatin plasticity represents a previously unrecognized mechanism linking neuronal metabolism to memory storage and learning-induced transcriptional events.

Here, we provide evidence demonstrating the role of Kcr as an epigenetic rheostat for hippocampal memory storage and glutamatergic neurotransmission. We show that spatial learning in mice increases Kcr levels in the dorsal hippocampus during the early temporal window of memory consolidation. Next, we demonstrate that depleting histone Kcr levels in the hippocampus by overexpressing the crotonyl-CoA hydratase CDYL leads to long-term spatial memory impairments. Conversely, increasing histone Kcr levels by oral administration of crotonate, a precursor of crotonyl-CoA, enhances hippocampus-dependent long-term memory. Using single-nuclei transcriptomic and chromatin accessibility analyses, we define subregion-specific molecular signatures of Kcr regulation across the hippocampus— revealing that Kcr enhancement selectively impacts genes encoding key regulators of glutamatergic neurotransmission and identifying enriched transcription factor motifs at their promoters that may govern their expression. We examine functional consequences of increasing histone Kcr by performing *ex vivo* real-time fluorescent imaging using genetically encoded glutamate and calcium sensors— demonstrating that Kcr enhancement augments presynaptic glutamate release and neuronal activity. Together, our findings reveal, for the first time, the role of a non-acetyl acyl modification in long-term memory and excitatory neurotransmission. More importantly, our study offers mechanistic evidence of how epigenetic tuning of transcriptional programs by Kcr translates into changes in neuronal function and behavioral outcomes—linking epigenetic events in the nucleus to synaptic activity.

## Results

### Spatial training induces lysine crotonylation in discrete hippocampal subregions

KATs catalyze the addition of crotonyl groups on histone lysine residues using crotonyl-CoA as a cofactor, and crotonyl-CoA levels are regulated by the enzymes ACSS2 and CDYL (**Fig. 1a**). We first investigated whether spatial learning induces changes in histone Kcr in the hippocampus, similar to the well-established increases in histone Kac following spatial training^2,9,34^. We trained wild-type (WT) mice in a spatial object recognition (SOR) task and examined Kcr levels in core histones extracted from the dorsal hippocampus at different time-intervals (0.5-, 1-, and 2-hrs) post training (**Supplementary Fig. 1a-b**). We focused on the dorsal hippocampus in all our experiments, given extensive evidence for its critical role in spatial memory consolidation^30,35-37^. Moreover, in the absence of prior knowledge regarding specific crotonylated histone lysine residues that respond to learning, we measured pan-lysine crotonylation (Kcr) to assess the global landscape of histone Kcr during spatial memory consolidation. Using an antibody that has been shown to selectively detect crotonylated lysine residues^12^ and has been extensively used to characterize Kcr in several other studies^24,38-41^, we observed that histone Kcr levels significantly increased at the 1-hr timepoint after SOR training and returned to baseline levels at the 2-hr timepoint (**Supplementary Fig. a-b**). Next, we performed immunofluorescence using the pan-Kcr antibody to map the spatial profiles of Kcr induction in the hippocampus following SOR training (**Fig. 1b-d**). Quantitative analysis of the nuclear Kcr levels revealed significant increases in the CA1 pyramidal layer and the upper blade of dentate gyrus (DG) (**Fig. 1c-d**), and in the subiculum (**Supplementary Fig. 2a-b**) at the 1-hr timepoint after training in the SOR task, whereas no significant changes in the nuclear Kcr levels were observed in hippocampal subregions CA3 and the lower blade of DG (**Fig. 1c-d**). These results identify histone Kcr to exhibit distinct spatiotemporal dynamics in the dorsal hippocampus following spatial learning.

**Figure 1.**
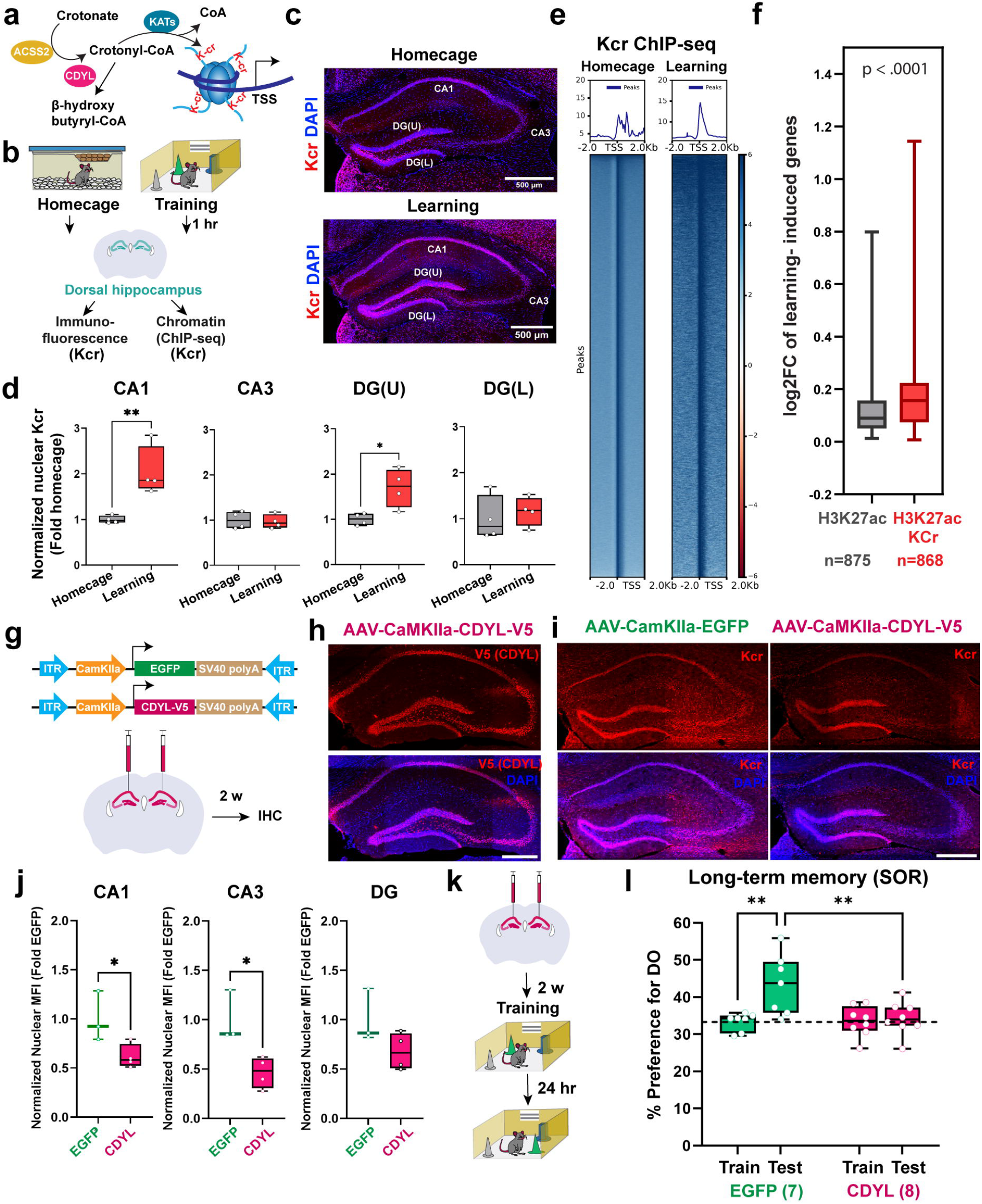
Lysine crotonylation is a critical regulator of learning-induced gene expression and long-term spatial memory consolidation. **a**. Depiction of molecular regulators of histone lysine crotonylation (Kcr). **b**. Schematic of the experiments performed in **c-i. c**. Immunofluorescence using anti-Kcr antibody showing levels of Kcr in different hippocampal sub-regions of HC and learning (SOR+1hr) mice. **d**. Normalized Mean Fluorescent Intensity (MFI) of nuclear Kcr levels across the groups. For CA1: unpaired t test: t (6)□= 3.810, **p□=□0.0089. Homecage (n□=□4) and SOR+1 hr (n□=□4), males only. For CA3: unpaired t test: t (6) = 0.2749, p□=□0.7926. Homecage (n□=□4) and SOR+1 hr (n□=□4), males only. For DG upper blade (U): unpaired t test: t (6)□= 3.118, *p□=□0.0206. Homecage (n□=□4) and SOR+1 hr (n□=□4), males only. For DG lower blade (L): unpaired t test: t (6)□= 0.5452, p□=□0.605. Homecage (n□=□4) and SOR+1 hr (n□=□4), males only. **e**. ChIP-seq density heat maps in dorsal hippocampus harvested from homecage (left) vs 1 hour after SOR-trained mice (right) for histone Kcr at ± 2 kb from the TSS. **f**. Box plots showing the extent of gene induction in the hippocampus following learning^28^ for genes that exhibit H3K27ac binding in the promoter region compared to genes that exhibit binding of H3K27ac and Kcr in the promoter region. Unpaired two-tailed t test was used to determine a significant change in mean log2FC between the groups. **g**. Schematic of viral constructs transduced in the dorsal hippocampus of adult male mice to overexpress Chromodomain Y-like protein (CDYL). AAV_9_-CaMKIIα-EGFP served as vector control, while AAV_9_-CaMKIIα-CDYL-V5 was used to drive expression of CDYL in excitatory neurons. **h**. Immunofluorescence using V5 antibody showing ectopic expression of CDYL-V5 in dorsal hippocampus two weeks after viral infusion. **i**. Immunofluorescence image of the dorsal hippocampus using pan-Kcr antibody in CDYL-V5 or control expressing mice. **j**. Normalized Mean Fluorescent Intensity (MFI) of nuclear Kcr levels across the groups. For CA1: Unpaired t test: t (5)□= 2.679, *p□=□0.439. EGFP (n□=□3) and CDYL (n□=□4), males only. For CA3: Unpaired t test: t (5) = 3.417, *p□=□0.0189. EGFP (n□=□3) and CDYL (n□=□4), males only. For DG: Unpaired t test: t (5)□= 1.863, p□=□0.1215. EGFP (n□=□3) and CDYL (n□=□4), males only. **k**. Long-term memory (24 hr) assessment of mice infused with AAV-EGFP or AAV-CDYL into dorsal hippocampus. **l**. Quantification of **k**. Two-way ANOVA: significant session x virus interaction F_(1,13)_=6.323, p=0.0259, Sidak’s multiple comparison tests, **p=0.0041 (EGFP 24 hr Test vs CDYL 24 hr Test) and **p=0.0051 (EGFP Training vs EGFP 24 hr Test); AAV-EGFP (n=7), AAV-CDYL (n=8), males only. All box plots: the center line represents the median, the box edges represent the top and bottom quartiles (25th to 75th percentiles), and the minimum and maximum whiskers.

### Enrichment of crotonylation on gene promoters correlates with higher magnitude of learning-induced gene expression

To determine whether Kcr has a role in regulating learning-induced gene expression, we performed Chromatin Immunoprecipitation followed by sequencing (ChIP-Seq) using a pan-Kcr antibody on dorsal hippocampal tissue from mice that were either housed in their homecage or trained in the SOR task (**Fig. 1b, e**). We found that spatial learning led to a significant enrichment of Kcr on gene promoters (**Fig. 1e**), consistent with previous reports linking the presence of Kcr marks on gene promoters to transcriptional activation. Next, we determined the relationship between Kcr and a transcriptionally active Kac mark (H3K27ac) on learning-induced gene promoters. We compared the enrichment of Kcr and H3K27ac^42^ on the promoters of genes upregulated after learning in the dorsal hippocampus following the SOR task^30^. Our analyses revealed that learning-induced genes enriched with H3K27ac and pan-Kcr exhibit a significantly higher magnitude of expression (fold change) compared to genes enriched with only H3K27ac **(Fig. 1f)**. These findings not only underscore the transcriptional activation potential of Kcr but also reveal its role in regulating learning-dependent gene expression.

### Reduction of hippocampal Kcr levels impairs long-term spatial memory

The crotonyl-CoA hydratase Chromodomain Y-like protein (CDYL) hydrolyzes crotonyl-CoA to beta-hydroxy butyryl-CoA (**Fig. 1a**), thereby reducing histone Kcr levels^24^. To date, CDYL remains the only bona fide negative regulator of Kcr that does not affect other histone acylations^39^—presenting us with a unique strategy to selectively manipulate histone Kcr levels and examine its impact on memory consolidation. Interestingly, neuronal activity has been shown to reduce CDYL levels^43^, prompting us to investigate whether spatial learning would similarly impact hippocampal CDYL levels. We performed immunofluorescence on brain sections obtained from SOR-trained WT mice to examine spatial patterns of CDYL protein levels following learning across hippocampal subregions (**Supplementary Fig. 3a-d**). Hippocampal subregions CA1, subiculum, and the upper blade of DG exhibited significant downregulation of nuclear CDYL levels 1-hr after SOR training, whereas nuclear CDYL levels remained unchanged in the hippocampal subregions CA3 and the lower blade of DG (**Supplementary Fig. 3a-d)**. The downregulation of nuclear CDYL levels after learning in CA1, subiculum, and the upper blade of DG offers a molecular explanation for the corresponding learning-induced increase in nuclear Kcr levels observed within these hippocampal subregions.

Next, to determine whether histone Kcr has a role in hippocampal long-term memory, we designed a genetic strategy to selectively modulate Kcr in the dorsal hippocampus by manipulating levels of CDYL. We injected an adeno-associated virus (AAV) in WT mice to overexpress CDYL (AAV_9_-CaMKIIα-CDYL-V5) in excitatory neurons of the dorsal hippocampus (**Fig. 1g**). Immunofluorescence performed two weeks after viral infusion confirmed the expression of CDYL-V5 across the dorsal hippocampal principal neuronal layers (**Fig. 1h**). We observed significantly reduced Kcr levels in the hippocampal areas CA1 and CA3 in CDYL-V5-expressing mice compared to EGFP controls (**Fig. 1i-j**), thus validating the efficacy of our viral-based approach. Next, we evaluated long-term spatial memory in mice infused with AAV-CDYL-V5 in the dorsal hippocampus. Mice were trained in the SOR task with objects that were placed in specific spatial locations within a SOR arena. To test for long-term spatial memory, we performed a retrieval task 24-hrs later with one of the objects displaced to a novel spatial location (**Fig. 1k**). Control AAV-infused mice (AAV_9_-CaMKIIα-EGFP) exhibited a higher preference towards the displaced object during the test session, suggesting intact memory consolidation.

Conversely, CDYL-infused mice (AAV_9_-CaMKIIα-CDYL-V5) failed to exhibit any preference towards the displaced object during the test session (**Fig. 1l**). Collectively, these findings suggest a critical role of hippocampal Kcr in spatial memory consolidation.

### Pharmacologically increasing Kcr enhances long-term memory

Given that a reduction in hippocampal Kcr levels significantly impairs long-term memory, we next asked whether increasing Kcr levels could enhance long-term memory. Administration of crotonate, a short-chain fatty acid, has been shown to elevate histone Kcr^12,23^ and stimulate gene expression^23^ by increasing Crotonyl-CoA levels^12,23^. Therefore, we implemented this strategy to pharmacologically increase histone Kcr levels in mice. We administered two different doses of crotonate (50 and 200 mg/kg) to WT mice by oral gavage and examined histone Kcr levels in the dorsal hippocampus 1-hr after drug administration (**Fig. 2a**). We observed a significant increase in histone Kcr following 200 mg/kg dose of crotonate, whereas the 50 mg/kg dose of crotonate failed to enhance histone Kcr levels compared to vehicle (saline)-treated mice (**Fig. 2b**). Next, to study the impact of higher histone Kcr levels on long-term memory, we trained WT mice in a weak-learning (sub-threshold) SOR paradigm^44^ to evaluate long-term memory enhancement. The subthreshold variant of the SOR task was designed to assess enhancements in long-term memory, as weak SOR training protocols typically do not result in long-term memory^44-46^. Mice were administered 50mg/kg of crotonate, 200 mg/kg of crotonate, or vehicle via oral gavage immediately after the completion of the training session. When tested 24 hrs later, both the vehicle-treated and 50 mg/kg crotonate-treated mice did not show any preference towards the displaced object, whereas mice administered with 200 mg/kg crotonate showed a significant preference towards the displaced object (**Fig. 2c**). Consistent with our SOR findings, we observed a significantly enhanced freezing response upon crotonate administration when mice were tested 24 hrs following a sub-threshold contextual fear conditioning (CFC) task (**Fig. 2d**). Our behavioral findings demonstrate that increases in histone Kcr augments hippocampal long-term spatial and contextual fear memories, further underscoring the functional relevance of Kcr in long-term memory consolidation.

**Figure 2.**
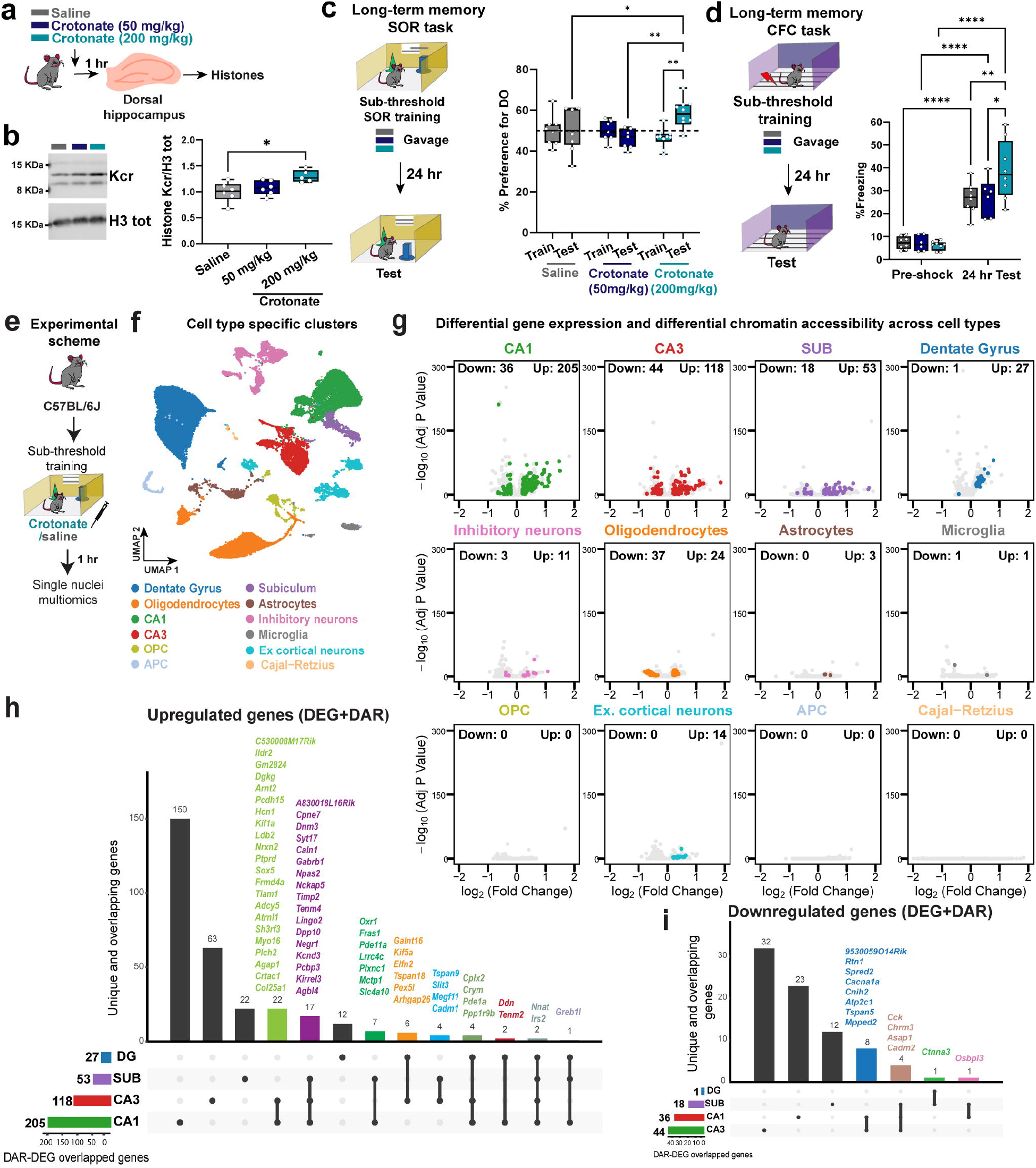
Single-nuclei multiomics (RNA+ATAC-seq) reveals mechanistic insights underlying lysine crotonylation-mediated memory enhancement. **a**. Kcr Western blots of core histones extracted from dorsal hippocampus 1 hr after crotonate treatment (50 mg/kg or 200 mg/kg dose). Vehicle (saline) treated mice were used as controls. **b**. Quantification of **a**. One-way ANOVA: Kcr: F _(2, 14)_ = 5.550, p=0.0168. Dunnett’s multiple comparisons tests: *p= 0.0108 (saline versus crotonate 200mg/kg). Saline (n=6), Crotonate 50 mg/kg (n=6), Crotonate 200 mg/kg (n=5), males only. **c**. Long-term memory enhancement in a sub-threshold SOR learning paradigm following crotonate treatment. Two-way ANOVA: significant session x treatment interaction F_(2, 23)_ = 5.081, p= 0.0149, Sidak’s post hoc tests, *p=0.0359 (200mg/kg crotonate 24 hr test vs saline 24 hr test), **p=0.0021 (200mg/kg crotonate train vs 200mg/kg crotonate 24 hr test), **p=0.0025 (200mg/kg crotonate 24 hr test vs 50mg/kg crotonate 24 hr test). Saline (n=9), Crotonate 50 mg/kg (n=7), Crotonate 200 mg/kg (n=10), males only. **d**. Long-term memory in a sub-threshold CFC learning paradigm following crotonate treatment. Two-way ANOVA: Significant session x treatment interaction F _(2, 19)_ = 4.946, *p=0.0187, main effect of session (pre-shock and 24hr test): F_(1, 19)_ = 140.9, p<0.0001. Sidak’s post hoc tests, *p=0.0130 (200mg/kg crotonate 24 hr test vs 50mg/kg crotonate 24 hr test), **p=0.0051 (200mg/kg crotonate 24 hr test vs saline 24hr test). ****p<0.0001 (Saline pre-shock vs saline 24hr test), ****p<0.0001 (crotonate 50mg/kg pre-shock vs crotonate 50mg/kg 24hr test), and ****p<0.0001 (crotonate 200mg/kg pre-shock vs crotonate 200mg/kg 24hr test). Saline (n=8), Crotonate 50 mg/kg (n=6), Crotonate 200 mg/kg (n=8), males only. **e**. Experimental scheme: Adult male C57BL/6J mice were trained in a sub-threshold SOR paradigm and administered with crotonate (200 mg/kg, oral gavage; n=4) or saline (oral gavage; n=4) immediately after the completion of training. 1-hr later, the dorsal hippocampus was harvested, and nuclei were isolated for single-nuclei multiomics processing. Hippocampi from two animals within the same group were pooled for each droplet capture, resulting in a final sample size of n=2 per group for the single-nuclei multiomics analysis. **f**. UMAP plot showing cell type-specific clusters of the dorsal hippocampus. **g**. Volcano plots depicting genes that exhibit differential expression and chromatin accessibility following crotonate treatment. Genes labeled with color are DEGs that also exhibit DARs, while depict genes that just exhibit differential expression but do not exhibit DARs are labeled in gray. **h-i**. Upset plots depicting unique and overlapping signatures of DEGs with DARs across all the hippocampal principal neurons. **h**. Upregulated DEGs with DARs. **i**. Downregulated DEGs with DARs. All box plots: the center line represents the median, the box edges represent the top and bottom quartiles (25th to 75th percentiles), and the minimum and maximum whiskers.

### Neuronal ACSS2 is a key regulator of Kcr-dependent memory enhancement

The metabolic enzyme ACSS2 has been shown to synthesize acetyl-CoA. Previous work has demonstrated that ACSS2 plays a critical role in long-term memory by coupling acetyl-CoA synthesis to Kac-mediated transcription of learning-responsive genes^33^. However, ACSS2 can also synthesize crotonyl-CoA from the short-chain fatty acid crotonate in mammalian cells^12^ (**Fig. 1a, Supplementary Fig. 4a**). This prompted us to investigate whether ACSS2 regulates the increases in histone Kcr and enhancements in long-term memory seen after the administration of crotonate. We injected an AAV vector (AAV_9_-CaMKIIα-Cre-EGFP) into the dorsal hippocampus of male ACSS2^f/f^ mice^47^ to achieve a conditional knockdown of ACSS2 expression selectively in the excitatory neurons of the dorsal hippocampus (**Supplementary Fig. 4b**). Immunofluorescence performed on brain sections obtained from the ACSS2 conditional knockout mice (ACSS2 cKO mice; ACSS2 ^f/f^:CaMKIIα-Cre-EGFP) revealed the expression of Cre across hippocampal subregions (**Supplementary Fig. 4c**). Western blot analyses confirmed the downregulation of ACSS2 in the dorsal hippocampus of ACSS2 cKO mice (**Supplementary Fig. 4d-e**). We then examined the impact of crotonate administration on histone Kcr levels in ACSS2 cKO mice. We administered crotonate (200 mg/kg) or vehicle (saline) via oral gavage immediately after a sub-threshold SOR training session, and quantified histone Kcr levels in the dorsal hippocampus 1-hr after training. We found that crotonate treatment failed to enhance hippocampal histone Kcr levels in ACSS2 cKO mice (**Supplementary Fig. 4f-g**), despite administering a dose of crotonate (200 mg/kg) that was sufficient to induce histone Kcr levels in WT mice (**Fig. 2b**). As crotonate administration failed to enhance Kcr levels in ACSS2 cKO mice, we hypothesized that crotonate treatment would also fail to elicit long-term memory enhancement in ACSS2 cKO mice. To test this hypothesis, we trained ACSS2 cKO mice in a sub-threshold learning SOR paradigm, administered vehicle or crotonate (200 mg/kg) immediately after training, and examined their long-term memory 24 hrs after training (**Supplementary Fig. 4h**). Our behavioral assessment confirmed that crotonate treatment in ACSS2 cKO mice had no impact in long-term memory compared to the vehicle-treated group (**Supplementary Fig. 4i**). Taken together, these results demonstrate that ACSS2 is critical in mediating the molecular impact of crotonate-mediated increases in Kcr on memory consolidation.

### Single-nuclei multiomics reveals the molecular signatures of Kcr underlying memory enhancement

Next, we implemented a single-nuclei multiomics (snRNA-seq and snATAC-seq) strategy to elucidate the molecular mechanisms underlying crotonate-mediated memory enhancement. Mice were trained in SOR using a sub-threshold learning paradigm, and oral administration of crotonate (200 mg/kg) or vehicle (saline) was performed immediately after training. 1-hr after crotonate administration, the dorsal hippocampus was harvested, and nuclei were isolated for single-nuclei multiomics (**Fig. 2e**). Cell type clustering identified well-distinguished clusters comprising excitatory and inhibitory neurons, as well as clusters representing non-neuronal cell populations (**Fig. 2f**). Excitatory neurons were further classified into CA1, CA3, subiculum (Sub), dentate gyrus, and excitatory cortical neurons based on cell-type-specific marker gene expression (**Fig. 2f, Supplementary Fig. 5a**). Notably, crotonate and vehicle treatment groups exhibited the same number of cell-type-specific clusters and comparable proportion of cell-types (**Supplementary Fig. 5b-c, Supplemental table 1**). Next, we performed differential gene expression (DEG) utilizing the snRNA data along with differential accessibility regions (DARs) on the chromatin from the snATAC data of each cluster, comparing crotonate and vehicle treated groups. For downstream analysis, we used the DEGs (FDR<0.05, and absolute log_2_ fold change > 0.2) exhibiting concordant DARs (FDR<0.05, and absolute log_2_ fold change > 0.2) across their promoter and gene body (**Fig. 2g, Supplemental table 2**). We found that excitatory neurons in the CA1 hippocampal area showed the highest number of genes impacted by crotonate, with 205 upregulated genes showing increased chromatin accessibility and 36 downregulated genes showing reduced chromatin accessibility (**Fig. 2g**). Other cell types that showed a strong impact of crotonate on gene expression and chromatin accessibility were CA3 excitatory neurons (118 upregulated genes with increased chromatin accessibility and 44 downregulated genes with reduced chromatin accessibility), subiculum (53 upregulated genes with increased chromatin accessibility and 18 downregulated genes with reduced chromatin accessibility), and the DG (27 upregulated genes with increased chromatin accessibility and one downregulated gene with reduced chromatin accessibility) (**Fig. 2g**). In addition to the principal neuronal clusters (CA1, CA3, subiculum, and DG), our analysis identified discrete molecular changes in inhibitory neurons (11 upregulated genes with increased DARs and 3 downregulated genes with reduced DARs) (**Fig. 2g**). Among the non-neuronal cells, oligodendrocytes showed the highest number of overlapping DEGs and DARs (24 upregulated and 27 downregulated) (**Fig. 2g**). Notably, we found a significant correlation between the significant differentially expressed genes and genes with significant differential chromatin accessibility in hippocampal subregions CA1 and CA3 (**Supplementary Fig. 6**). Next, utilizing an Upset plot, we compared unique and overlapping upregulated genes with concordant DARs in each cell-type. We found that CA1, CA3, subiculum, and DG showed a common upregulation of 2 genes (**Fig. 2h**). Notably, CA1, CA3, and subiculum showed 17 commonly upregulated genes, CA1, CA3, and DG showed a common upregulation of 4 genes, and CA1 and CA3 showed 22 commonly upregulated genes following crotonate treatment (**Fig. 2h**). Like the cell-type expression patterns observed with upregulated DEGs with increased DARs, there was some overlap in the expression of downregulated genes with reduced DARs between cell-types. We identified 23 downregulated genes specific to CA1, 32 exclusively in CA3, and 12 selectively in the subiculum (**Fig. 2i**). Four genes were commonly downregulated across CA1, CA3, and the subiculum, while 8 downregulated genes were shared between CA1 and CA3 (**Fig. 2i**). Because the manipulation of Kcr levels selectively in excitatory neurons was found to impact long-term memory (**Fig. 1g-l, Supplementary Fig. 4**), our subsequent analyses focused on DEGs within the neuronal populations of CA1, CA3, subiculum, and DG that also exhibit DARs.

To better understand the impact of Kcr enhancement on the molecular signatures of principal neuronal sub-types in the dorsal hippocampus, we performed Gene Ontology (GO: Molecular Function) overrepresentation analysis to identify the pathways enriched among the DEGs with DARs following crotonate treatment. We found that crotonate treatment impacted the expression of genes that were predominantly associated with regulating neurotransmission and synaptic function across the cell clusters (**Fig. 3a-d)**. Among the most significant pathways found enriched following crotonate administration, cell adhesion molecule binding was commonly enriched in subregions CA1, CA3, and subiculum, whereas distinct pathways attributed to ion channel activity were commonly enriched in CA1 and CA3 (**Fig. 3a-d, Supplemental table 3**). In contrast, pathways attributed to PDZ domain binding and calcium-dependent phospholipid binding were found enriched in the subiculum, whereas DG exhibited an enrichment of pathways related to protein kinase C activity and serine/threonine kinase activity. (**Fig. 3a-d**). Among the downregulated DEGs with reduced chromatin accessibility, we found pathways attributed to ephrin receptor activity, calcium ion transmembrane transporter activity, and phosphoric diester hydrolase activity downregulated in CA3, whereas pathways involved in cadherin binding and beta-catenin binding were downregulated in DG (**Supplementary Fig. 7**).

**Figure 3.**
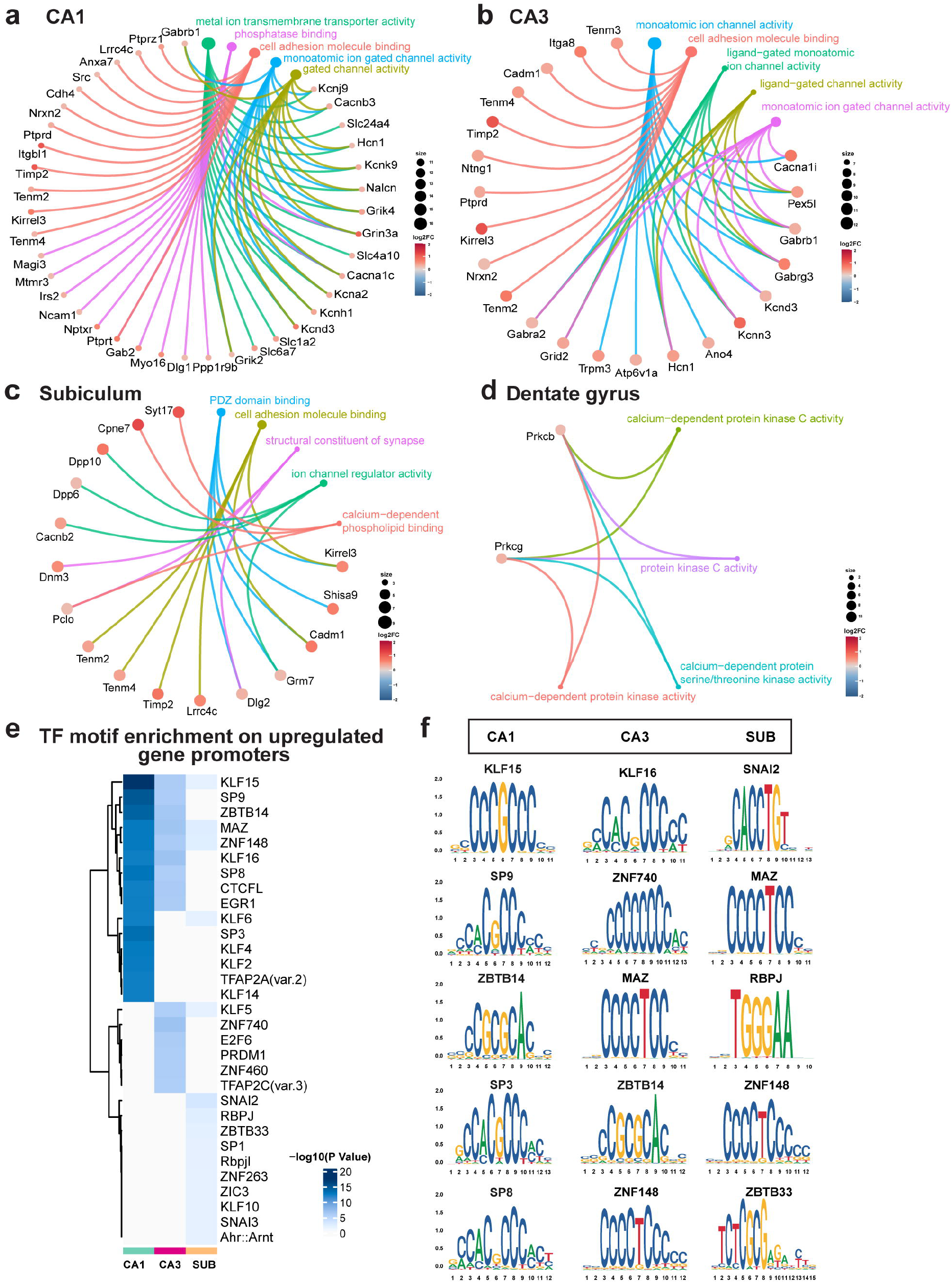
Increases in Kcr elicits distinct molecular signatures in principal hippocampal neuronal cell-types. **a-d**. Cnet plots depicting the top five enriched pathways (GO: Molecular Function) and their corresponding DEGs with DARs in hippocampal subregions CA1 **(a)**, CA3 **(b)**, Subiculum **(c)**, and Dentate gyrus **(d)** that are upregulated in response to crotonate administration. **e-f**. Single-nuclei multiomics reveals transcription factor motif sites enriched on gene promoters that show differential expression and differential chromatin plasticity following crotonate treatment. **e**. Transcription factor (TF) motif enrichment analysis from upregulated genes with increased chromatin accessibility at the promoter upon crotonate administration. **f**. Top 5 significant TF motifs enriched from CA1, CA3, and subiculum.

To gain a mechanistic understanding of how crotonate-mediated changes in chromatin accessibility activate gene transcription, we performed a Transcription Factor (TF) Motif Enrichment analysis on the differentially accessible promoter regions of the upregulated DEGs (**Fig. 3e-f, Supplemental table 4**). Among the top 15 most significant TF motifs enriched on the promoters of upregulated DEGs across the cell clusters, TF motifs KLF15, ZNF148, and MAZ were found commonly enriched across subregions CA1, CA3, and subiculum. Commonalities in the enrichment of TF motifs on upregulated DEG promoters were observed across hippocampal subregions, such as EGR1 and CTCFL in CA1 and CA3. No significant TF motifs were found enriched in the upregulated DEGs in the DG.

### Cell-cell communication analysis reveals enhanced glutamate signaling following increases in Kcr

Activity-dependent synaptic transmission within the intrahippocampal circuit facilitates adaptive behaviors for encoding spatial, episodic, and contextual memories^48-50^. Utilizing published databases of ligand-receptor interactions^51^, we constructed an intercellular communication network of the principal neuronal subtypes to study the impact of Kcr enhancement on intrahippocampal signaling. Our analysis revealed that crotonate administration primarily increases the strength of glutamatergic signaling between the principal neuronal layers within the hippocampal circuit (**Fig. 4a**). Next, we investigated the ligand-receptor pairs in each intrahippocampal connection that exhibit increase in their communication probability following crotonate administration. We found that increasing Kcr levels strengthened the communication between distinct glutamatergic ligand-receptor pairs within specific neuronal networks of the hippocampal circuit. (**Fig. 4b**). These include the Slc1a2-Grik2/Grik4/Grik5, and Slc1a2-Gria4 pairs in the DG-CA3, Slc1a2-Grik2/Grik5/Grm7, and Slc1a1-Grik2/Grik5/Grm7 pairs in the CA3-CA1, and Slc1a1-Grik2/Grik5/Grm7, Slc1a2-Gria3/Gria4/Grik2/Grik5, and Slc17a7-Grik2/Gria3/Grm7/Grm8 pairs in the CA1-subiculum neuronal connection (**Fig. 4b**). We computed the communication probability of ligand-receptor interactions for glutamate signaling genes within principal neuronal cell types, which further confirmed the enhanced strength of glutamatergic transmission after crotonate administration (**Fig. 4c**). Collectively, our findings suggest that crotonate-dependent increase in histone Kcr augments the strength of glutamatergic neurotransmission within the hippocampal circuit.

**Figure 4.**
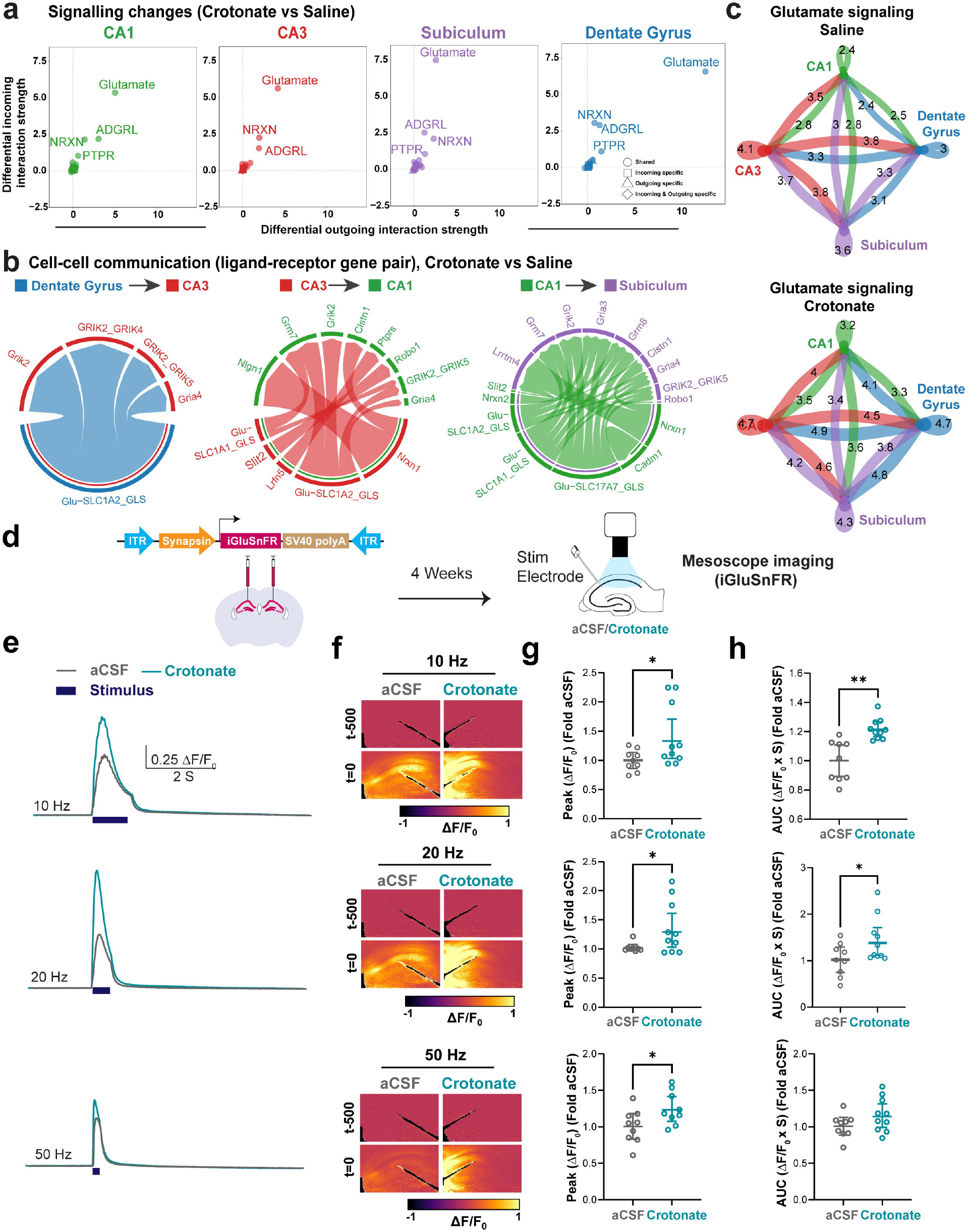
Increases in Kcr augments glutamatergic neurotransmission. ***a***. Differential incoming interaction strength and differential outgoing interaction strength in crotonate-treated mice compared to saline-treated mice in hippocampal subregions CA1, CA3, Subiculum, and DG. Differential changes in interaction strength in ‘Shared’ (both incoming and outgoing connections in both the groups), ‘Incoming specific’ (incoming connections in both the groups), ‘Outgoing specific’ (outgoing connections in both the groups), and ‘Incoming & Outgoing specific’ (both incoming and outgoing connections in either of the groups) clusters are depicted by symbols. **b**. Individual ligand-receptor interactions significantly enhanced upon crotonate treatment in intrahippocampal projections of DG-CA3, CA3-CA1, and CA1-subiculum. **c**. Interaction strength of ligand-receptor genes involved in glutamatergic signaling across the principal hippocampal neuronal networks in saline-treated and crotonate-treated mice. Communication probability, represented by numbers ranging from 0 to 10, is visualized through the edge width of intrahippocampal connections, indicating the strength of communication between the respective subregions. **d-h**. Kcr enhancement leads to larger stimulus-evoked glutamate transients in acute hippocampal slices. **d**. Schematic of the experimental design. **e**. Representative traces of evoked glutamate transient responses at 10, 20 and 50 Hz in control slices and ones incubated with crotonate (5mM). **f**. Image panels showing stimulus-evoked glutamate transient response over time in the hippocampus upon stimulating the Schaffer collaterals in the CA1 Stratum Radiatum. The black object in each image depicts the stimulating electrode. t=0 denotes the time point corresponding to the peak amplitude; t−500 represents the time point 500 ms prior to t=0. **g**. The peak amplitude of evoked glutamate transients was significantly higher in crotonate-treated slices compared to control across all the stimulation frequencies. Normalized Peak Amplitude across the groups. For 10 Hz (top): Lognormal Welch’s t test: t (13.96)□= 2.37, *p□=□0.0327. aCSF (n□=□9) and Crotonate (n□=□10). For 20 Hz (middle): Lognormal Welch’s t test: t (9.982) = 2.302, *p□=□0.0442. aCSF (n□=□9) and Crotonate (n□=□10). For 50 Hz (bottom): Lognormal Welch’s t test: t (14.72) = 2.274, *p□=□0.0384. aCSF (n□=□9) and Crotonate (n□=□10). **h**. The area under the curve (AUC) of evoked glutamate transients was significantly higher in crotonate-treated slices compared to control in the 10 Hz and 20 Hz stimulation groups. For 10 Hz (top): Lognormal Welch’s t test: t (10.34)□= 4.043, **p□=□0.0022. aCSF (n□=□9) and Crotonate (n□=□10). For 20 Hz (middle): Lognormal Welch’s t test: t (14.96) = 2.263, *p□=□0.039. aCSF (n□=□9) and Crotonate (n□=□10). For 50 Hz (bottom): Lognormal Welch’s t test: t (16.92) = 1.625, p□=□0.1226. aCSF (n□=□9) and Crotonate (n□=□10). All scatter plots: the center line represents the mean, error bars represent ±SEM.

### Real-time ex vivo fluorescent imaging reveals enhanced activity-dependent glutamate transients following increases in Kcr

Our single-nuclei multiomics and cell–cell communication analyses indicated enhanced glutamatergic signaling—a functional readout of which is presynaptic glutamate release. To test this, we transduced WT mice with an AAV encoding the fluorescent glutamate sensor iGluSnFR (pAAV_9_-hSyn-iGluSnFR) in the dorsal hippocampus. Four weeks later, acute hippocampal slices were prepared and incubated in either artificial CSF (aCSF) or aCSF containing 5 mM crotonate (**Fig. 4d**). We applied three stimulation frequencies (10, 20, and 50 Hz) based on prior studies^52^ and recorded stimulus-evoked glutamate transients in CA1 following stimulation of CA3–CA1 Schaffer collateral afferents in the stratum radiatum (**Fig. 4e–h**). Crotonate-treated slices exhibited a significant increase in peak glutamate amplitude across all three stimulation frequencies (**Fig. 4g**), with a concomitant increase in area under the curve (AUC) at 10 and 20 Hz stimulation (**Fig. 4h**). In contrast, glutamate clearance kinetics (□_fast_ and □_slow_; depicted in a representative trace illustrated in **Supplementary Fig. 8**) were unchanged across conditions (**Supplementary Fig. 9a-b**), suggesting that there is no difference in glutamate clearance. Together, these results indicate that Kcr augments glutamatergic neurotransmission by facilitating activity-dependent presynaptic glutamate release.

### Real-time ex vivo fluorescent imaging reveals enhanced post-synaptic responsiveness to Kcr-dependent increase in presynaptic glutamate release

Our iGluSnFR findings demonstrating increased presynaptic glutamate release complements our cell– cell communication analyses, altogether indicating enhanced glutamatergic signaling strength with elevated Kcr. This led us to hypothesize that Kcr-mediated enhancement of activity-dependent presynaptic glutamate release at CA3–CA1 synapses would alter circuit responsiveness to increased glutamatergic input. Such changes in circuit activity could be mediated, in part, by select postsynaptic glutamate receptor genes that regulate activity-dependent calcium influx—the expression of which were found upregulated following crotonate administration. As a proof of concept, we investigated the expression profile of the AMPA receptor encoding gene *Gria4*^*53*^, one of the significantly upregulated glutamatergic signaling genes following crotonate treatment. Consistent with the single-nuclei transcriptomic data (**Fig. 5a**), RNAscope analysis revealed a significant increase in *Gria4* expression in hippocampal CA1 after crotonate administration (**Fig. 5b-c**).

**Figure 5.**
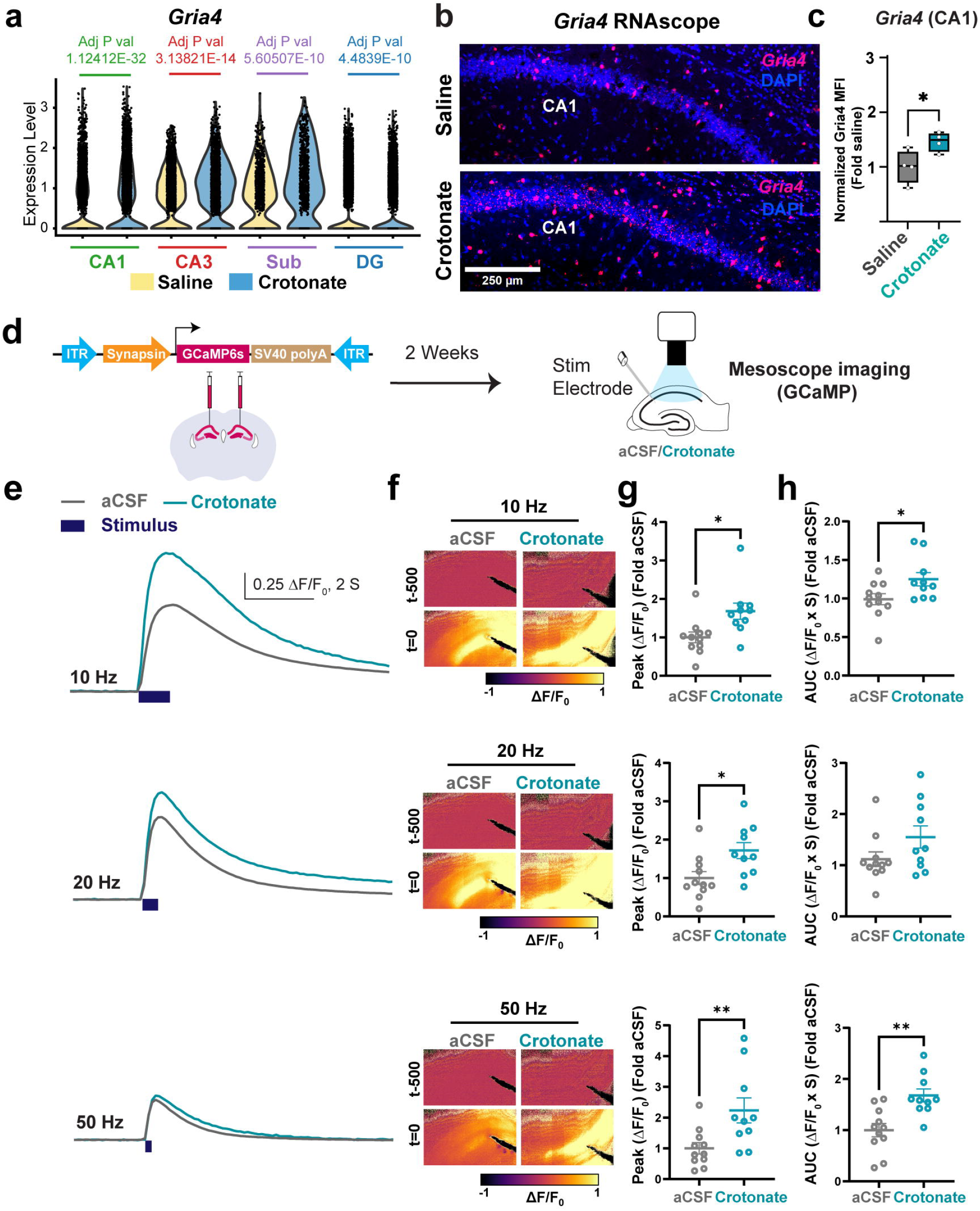
Kcr enhancement amplifies post-synaptic responsiveness to increases in presynaptic glutamate release. **a**. Violin plot depicting expression of *Gria4* mRNA in hippocampal principal neuronal cell types across saline- and crotonate-treated conditions. **b**. RNAscope validation of crotonate-mediated increases in *Gria4* expression in hippocampal CA1. Mice were trained with a sub-optimal SOR training protocol, administered saline or crotonate (200 mg/kg) immediately after training, and perfused 1 hr later. Scalebar: 100 µm. **c**. Quantification of **b**. For CA1: Normalized Mean Fluorescent Intensity (MFI) of *Gria4* levels across the groups. Unpaired t test: t (6)□= 2.613, *p□=□0.04. Homecage (n□=□4) and SOR+1 hr (n□=□4), males only. **d-h**. Kcr enhancement leads to larger stimulus-evoked calcium transients in acute hippocampal slices. **d**. Schematic of the experimental design. **e**. Representative traces of evoked calcium transient responses at 10, 20 and 50 Hz frequencies in control and crotonate (5mM)-incubated slices. **f**. Image panels showing stimulus-evoked calcium transients over time in the hippocampus upon stimulating the Schaffer collaterals in the CA1 Stratum Radiatum. The black object in each image depicts the stimulating electrode. t=0 denotes the time point corresponding to the peak amplitude; t−500 represents the time point 500 ms prior to t=0. **g**. The peak amplitude of evoked calcium transients was significantly higher in crotonate-treated slices compared to control across all the stimulation frequencies. Normalized Peak Amplitude across the groups. For 10 Hz (top): Lognormal Welch’s t test: t (17.84)□= 2.803, *p□=□0.118. aCSF (n□=□11) and Crotonate (n□=□10). For 20 Hz (middle): Lognormal Welch’s t test: t (17.18) = 2.734, *p□=□0.014. aCSF (n□=□11) and Crotonate (n□=□10). For 50 Hz (bottom): Lognormal Welch’s t test: t (18.98) = 3.134, **p□=□0.0055. aCSF (n□=□11) and Crotonate (n□=□10). **h**. The area under the curve (AUC) of evoked calcium transients was significantly higher in crotonate-treated slices compared to control in the 10 Hz and 50 Hz stimulation groups. For 10 Hz (top): Lognormal Welch’s t test: t (18.06)□= 2.260, *p□=□0.0364. aCSF (n□=□11) and Crotonate (n□=□10). For 20 Hz (middle): Lognormal Welch’s t test: t (18.43) = 1.702, p□=□0.1056. aCSF (n□=□11) and Crotonate (n□=□10). For 50 Hz (bottom): Lognormal Welch’s t test: t (13.52) = 3.241, **p□=□0.0062. aCSF (n□=□11) and Crotonate (n□=□10). All scatter plots: the center line represents the mean, error bars represent ±SEM. All box plots: the center line represents the median, the box edges represent the top and bottom quartiles (25th to 75th percentiles), and the minimum and maximum whiskers.

We then employed an *ex vivo* fluorescence imaging approach using the genetically encoded calcium indicator GCaMP6s to examine widefield CA1 circuit dynamics in response to Kcr-dependent increase in activity-dependent presynaptic glutamate release from CA3 afferents (demonstrated earlier in **Fig. 4d-h**). Wild-type mice were transduced with an AAV encoding GCaMP6s (pAAV_9_-hSyn-GCaMP6s-P2A-mRuby3) in the dorsal hippocampus, and acute slices were prepared two weeks later. Slices were incubated in either crotonate (5mM) or aCSF (**Fig. 5d**), and stimulus-evoked calcium transients were recorded over time at three different frequencies (**Fig. 5e-f**). Crotonate-treated slices exhibited significantly higher peak calcium transients across all three stimulation frequencies (**Fig. 5g**), accompanied by increased AUC at 10 and 50 Hz (**Fig. 5h**). To confirm that these effects were driven by transcriptional regulation rather than non-genomic actions of the drug, we performed a control experiment in which slices were incubated with crotonate for only 10 minutes (**Fig. S10**). This brief exposure failed to alter either peak amplitude or AUC (**Supplementary Fig. 10b-c**). Collectively, these findings indicate that elevated histone Kcr enhances activity-dependent calcium signaling in hippocampal neurons—an effect likely attributable to the transcriptional upregulation of postsynaptic calcium-permeable glutamate receptor genes following Kcr elevation.

## Discussion

The discovery of various non-acetyl histone acylations over the past decade has led to studies to determine whether these modifications play a unique role in the epigenetic regulation of gene expression. Histone Kac has long been regarded as a cornerstone of the epigenetic mechanisms underlying long-term memory consolidation^2,4,9,54^. In this study, we demonstrate the critical role of lysine crotonylation, a non-acetyl histone acylation associated with transcription activation^6,12,27^, in regulating hippocampal long-term memory consolidation. Through pharmacological and genetic approaches, we demonstrate the bidirectional modulation of long-term memory by manipulating Kcr levels in the dorsal hippocampus. Mechanistically, our findings reveal that memory enhancement observed upon pharmacologically increasing Kcr levels is mediated through the function of ACSS2 in excitatory neurons in the hippocampus. Single-nuclei transcriptomic and epigenomic analyses revealed that Kcr primarily regulates the chromatin accessibility and expression of genes linked to synaptic function and glutamatergic neurotransmission within principal neuronal populations of the dorsal hippocampus. We then performed cell–cell communication analyses which predicted that increasing hippocampal Kcr levels during the critical window of memory consolidation enhances the communication probability of glutamatergic neurotransmission within intrahippocampal circuits. We performed *ex vivo* real-time fluorescence-based imaging and found that Kcr indeed facilitates activity-dependent presynaptic glutamate release and enhances neuronal excitability. Altogether, our study identified a novel epigenetic mechanism critical for long-term memory storage and excitatory neurotransmission.

Histone acylation profiles are regulated not just by the enzymatic activities of their respective writers and erasers, but also through changes in the nuclear concentrations of their metabolic precursors^6,12^. We found that bidirectional manipulation of hippocampal crotonyl-CoA levels had contrasting effects on memory consolidation. Specifically, increasing crotonyl-CoA levels by administering crotonate increased histone Kcr levels in the dorsal hippocampus and enhanced long-term memory. Conversely, depletion of crotonyl-CoA levels by overexpressing CDYL reduced histone Kcr levels and led to deficits in long-term memory. Together, these findings not just establish Kcr as a key regulator of long-term memory but also identify CDYL as a potential memory suppressor gene^13^. Crotonyl-CoA levels are also regulated by the metabolic enzyme ACSS2, which synthesizes crotonyl-CoA from crotonate^12,32^. In our study, we found that crotonate fails to increase hippocampal histone Kcr levels or enhance long-term memory in the ACSS2 cKD mice, indicating that ACSS2 is essential for maintaining the nuclear pool of crotonyl-CoA. Collectively, these findings suggest that endogenous crotonate serves as a principal source of nuclear crotonyl-CoA, and that ACSS2-mediated conversion of crotonate to crotonyl-CoA is a critical molecular underpinning of Kcr-dependent enhancements in hippocampal memory.

Considering the promiscuity of histone acylation writers and erasers, the availability of metabolic precursors becomes even more critical in defining the landscape of histone acylations. Emerging evidence indicates that different acyl-CoA species compete for binding to KATs, such that changes in the relative abundance of specific acyl-CoAs can alter the balance of histone acylation marks^6,12^. For instance, depletion of cytoplasmic and nuclear acetyl-CoA pools in metazoans by knockdown of the enzyme ATP citrate lyase (ACLY) led to an increase in p300-catalyzed histone Kcr^55^. Thus, increasing the nuclear availability of less abundant acyl-CoA species enhances their access to shared acyltransferases, which in turn facilitates the catalysis of histone acylation marks. Canonical Kac writers such as p300, and readers such as the YEATS-domain proteins, catalyze and recognize Kcr more efficiently^27,56^, and Kcr itself serves as a potent activator of transcription^12^. Moreover, the thermal instability of the carbon–carbon double bond in crotonyl-CoA facilitates the dissociation of the crotonyl group from its CoA counterpart, thereby making it more convenient for the writers to efficiently deposit crotonyl moieties onto histone lysine residues^24,57^. Together, these observations raise the possibility that metabolically steering cells toward crotonyl-CoA utilization may represent an effective strategy to regulate transcriptional events during memory consolidation. In line with this idea, our findings show that pharmacologically increasing crotonyl-CoA levels to favor histone Kcr indeed offers a tractable and physiologically efficient approach to enhance long-term memory—laying the conceptual foundation to leverage Kcr as a “memory enhancer”.

Our single-nuclei multiomics approach, aimed at elucidating the molecular underpinnings of Kcr-mediated hippocampal memory enhancement, revealed unique signatures of differential gene expression and corresponding chromatin accessibility profiles within the hippocampal principal neurons. Surprisingly, most of the upregulated genes with increased chromatin accessibility following crotonate administration are not the canonical ‘Immediate Early Genes’ (IEGs) that serve as defining markers of engram ensembles activated by spatial learning^30,58-60^. Chromatin enrichment studies have shown that an increase in histone Kcr correlates with a reduction in the transcriptionally repressive mark (H3K27me3) and an increase in the transcription activation mark (H3K4me3) on gene promoters, with no observed change in the occupancy of acetylated histones (H3K27ac)^23^. Such an interplay between histone Kcr and other post-translational modifications, such as Kac and methylation, likely represents the framework of a unique “histone code” ^61-63^—the mechanistic framework of which remains to be elucidated in regard to long-term memory consolidation.

An important advantage of our multiomics framework is that it enabled us to interrogate the chromatin accessibility landscape and identify cell-type-specific signatures of TF motif enrichment that underlie Kcr-dependent gene regulatory programs. Notably, binding motifs of several members of the Krüppel-like family of transcription factors, such as KLF15 and ZNF148, were found enriched in hippocampal subregions CA1, CA3, and subiculum following Kcr enhancement. These TFs—integral in regulating diverse developmental and homeostatic functions—are increasingly being recognized for their role in long-term memory and plasticity. For instance, *Klf15*-null mice show marked deficits in hippocampal-dependent memory, including reduced contextual freezing after aversive learning^64^. Likewise, human genetic studies have shown that heterozygous variant in ZNF148 gene causes intellectual disability, whereas *de novo* mutations in ZNF148 have been associated with autism spectrum disorder, and cognitive impairment^65^. Beyond the Krüppel-like family, we also observed enrichment of MAZ binding across the gene bodies of Kcr-responsive hippocampal DEGs. MAZ is known to recognize a 27-bp GC-rich region within the *GluN1* promoter, and binding of MAZ to *GluN1* is essential for its transcriptional activation^66^. Another notable finding was the enrichment of EGR1 transcription factor motif in CA1 and CA3. *Egr1* (also known as *Zif268, Krox-24*, and *NGFI-A*) is a critical regulator of long-term memory and synaptic plasticity^67-70^. Beyond its canonical transcriptional regulatory function, recent studies have highlighted a link between Egr1 and epigenetic plasticity. Egr1 interacts with the DNA demethylase TET1 during postnatal brain development to program the brain methylome, with Egr1 binding sites becoming hypomethylated in mature neurons to regulate downstream gene expression^71^. Additionally, Egr1 knockdown increases histone acetylation at enhancer regions, regulating inflammatory gene expression in human macrophages^72^. Our findings extend the role of Egr1 in transcriptional regulation—indicating that it may play a previously unappreciated role in coordinating Kcr-mediated chromatin remodeling and gene expression. Collectively, these motif enrichments point to a regulatory architecture in which Kcr-mediated chromatin remodeling facilitates selective TF engagement with the genome, thereby regulating the transcriptional states that facilitate memory enhancement.

Another major strength of our study derives from our findings that increases in Kcr augments the dynamics of intrahippocampal glutamatergic neurotransmission. Our single-nuclei multiomics findings revealed that crotonate administration alters chromatin accessibility to upregulate genes involved in glutamatergic neurotransmission within excitatory neuronal populations. Cell-cell communication analysis—a powerful computational tool to infer and predict intercellular communication strength between neuronal populations within defined circuits^73^, further revealed that Kcr-enhancement augments glutamatergic neurotransmission within hippocampal circuit. Specifically, the analysis identified strengthened glutamatergic ligand–receptor interactions following Kcr enhancement, exemplified by an increased predicted communication probability between presynaptic vesicular glutamate transporters (VGLUTs) and postsynaptic kainite (KARs) and AMPA receptors (AMPARs). Together, these findings are suggestive of improved dynamics and fidelity of intrahippocampal glutamatergic signaling. These predictions were corroborated by *ex vivo* real time imaging, which demonstrated elevated activity-dependent presynaptic glutamate release in the CA3-CA1 Schaffer collaterals. This led to enhanced activity-dependent CA1 responsiveness at the circuit-level, as evidenced by amplified calcium transients *ex vivo* in response to increased presynaptic glutamate release at CA3–CA1 synapses. Importantly, studies examining histone PTMs on genes involved in glutamatergic neurotransmission are limited^74,75^, and, to date, no work has directly elucidated how epigenetic marks regulating glutamatergic gene expression bring about functional changes in synaptic neurotransmission and neuronal excitability. Our study reveals a mechanistic blueprint of how epigenetic regulation of glutamatergic neurotransmission shapes synaptic function and drives hippocampal memory consolidation.

In summary, our work demonstrates the critical role of a non-acetyl histone acylation that has previously not been studied in the context of long-term memory. We establish histone Kcr as a molecular switch for long-term memory, redefining our conceptual understanding of epigenetic mechanisms underlying long-term memory consolidation. We identify transcriptomic signatures of learning in distinct hippocampal subregions regulated by Kcr and highlight the epigenetic control of glutamatergic neurotransmission as a critical mechanism underlying Kcr-dependent long-term memory enhancement. By demonstrating that histone Kcr lies at the molecular crossroads of learning-induced transcriptional programs, synaptic neurotransmission, and behavior, our study provides a mechanistic framework for how epigenetic mechanisms translates into functional outcomes—shedding light on the elusive histone code of memory consolidation. Given that recent studies have demonstrated the role of Kcr-mediated regulatory mechanisms in several neurological disorders^39,76^, the broader relevance of our work is that it lays the groundwork to develop more effective therapeutic interventions to treat cognitive impairments associated with neurological, neurodevelopmental, and neuropsychiatric disorders.

## Methods

### Data reporting

No statistical methods were applied to predetermine sample size.

### Mouse lines

Adult male C57BL/6J mice were purchased from Jackson Laboratories. The ACSS2^f/f^ mice were generated at the University of Iowa Genome Editing Facility^47^. This mouse model was created by inserting loxP sites flanking the Exon2 of ACSS2 using CRISPR/Cas9. The mice were 2-4 months of age at the time all the behavioral and biochemical experiments were performed. All mice had ad libitum access to food and water, and lights were maintained on a 12-hr light/dark cycle. All experiments were conducted according to US National Institutes of Health guidelines for animal care and use and approved by the Institutional Animal Care and Use Committee of the University of Iowa, Iowa.

### Drug preparation

Sodium crotonate was freshly prepared on the day of the experiment. 2.5 gm of Crotonic acid (Sigma) was dissolved in 30 ml of 0.9% saline. Thereafter, the pH of the solution was adjusted to 7.4 using 5(N) NaOH, and additional 0.9% saline was added to adjust the final volume to 50 ml. The solution was then filtered through a 0.2 µm syringe filter. Appropriate volume of the drug (calculated based on the body weight of the animals) was administered via oral gavage based on the dosing regimen.

### Histone extraction

Histone extraction was performed using a commercially available kit (Histone Extraction Kit, Active Motif, #40028) according to the manufacturer’s protocol. Briefly, flash frozen hippocampi were mechanically homogenized in 300 μl of ice-cold Lysis Buffer AM8 using Dounce homogenizers and incubated on ice for 30 mins. The homogenate was centrifuged at 2,644 x g for 2 minutes at 4°C, and the nuclear pellet was resuspended in 250 μl ice-cold Extraction buffer. Following resuspension, the nuclear suspension was incubated on an end-to-end rotator overnight at 4°C. The pellet insoluble material was centrifuged at 20,800 x g for 10 minutes at 4°C the next day, the supernatant was collected, and the requisite volume of Complete Neutralization Buffer was added. Samples were stored in -80^°^C prior to western blotting.

### Western blot analysis

Western blotting was performed as previously described^30,58^. Whole cell lysates were run on a 4–20% Tris-HCl Protein Gel (BIO-RAD, #3450033). Proteins from the gel were then transferred to methanol-activated polyvinylidene difluoride membranes using the Trans-Blot Turbo Transfer System (BIO-RAD). Membranes were blocked with the Odyssey® Blocking Buffer (LI-COR) diluted in TBS and incubated overnight at 4°C with the following primary antibodies: pan-Kcr (1:1000, PTM BIO, #PTM-501), Total H3 (1:5000, ACTIVE MOTIF, #61647), ACSS2 (1:1000, Cell Signaling, #3658), and Actin (1:10,000, #MA1-91399). Post primary antibody incubation, membranes were washed thrice in TBS for 5 mins and incubated with IRDye® 800CW Goat anti-Rabbit IgG Secondary Antibody (1:5,000, LI-COR, #926-32211) and IRDye® 680LT Goat anti-Mouse IgG Secondary Antibody (1:5000, LI-COR, #926-68020). Membranes were then washed thrice in TBS, 5 mins each. Images were acquired using the Odyssey Infrared Imaging System (LI-COR). Quantification of western blot bands was performed using Image Studio Lite ver5.2 (LI-COR).

### Immunofluorescence

IHC experiments were performed as per earlier studies^30,58^. Animals were perfused with 4% PFA. Whole brains were harvested, immersed in 4% PFA, and stored at 4^°^C. 24 hrs after, brains were immersed in 30% sucrose and stored at 4^°^C. 20 μm coronal brain sections were made in a cryostat (Leica). Free-floating sections were washed thrice with PBS, blocked in a blocking buffer (0.1% PBST and 5% BSA) and incubated with the following primary antibodies for 48 hr: pan-Kcr (1:2000, PTM BIO, #PTM-501), CDYL (1:500, Sigma, #HPA035578), V5 (1:500, ThermoFisher Scientific, #37-7500), and GFP (1:4000, Abcam, #ab290). Post primary antibody incubation, sections were washed in PBS thrice, followed by a 2 hr secondary antibody incubation with the following antibodies: Goat anti-Mouse IgG (H+L) Cross-Adsorbed Secondary Antibody, Alexa Fluor™ 546 (1:500, ThermoFisher Scientific, #A-11003), Goat anti-Rabbit IgG (H+L) Cross-Adsorbed Secondary Antibody, Alexa Fluor™ 647 (1:500, ThermoFisher Scientific, #A-21244). Sections were then washed thrice in PBS and mounted on Superfrost™ Plus microscope slides (Fisherbrand). This was followed by coverslip mounting with ProLong™ Diamond Antifade Mountant with DAPI (ThermoFisher Scientific, #P36962).

### Confocal imaging and image analysis

Confocal images of IHC experiments were obtained in an Olympus FV 3000 confocal microscope using a 20X NA = 0.4 achromat dry objective at 800 × 800-pixel resolution and 1X optical zoom. All images (16 bit) were acquired with identical settings for laser power, detector gain and pinhole diameter for each experiment and between experiments. Images from the different channels were stacked and projected at maximum intensity using ImageJ (NIH). Mean fluorescence intensity (MFI) was computed using ImageJ plugins. MFIs were then subtracted to background intensities in each slice. Background subtracted MFIs in the experimental group were then normalized to the average background subtracted MFIs in control. For MFI quantification, the entire area corresponding to CA1, CA3, DG or subiculum were considered as region of interest.

### Chromatin Immunoprecipitation (ChIP)

Frozen hippocampal tissue samples (∼20 mg) was crosslinked in 1% formaldehyde for 10 min at room temperature with gentle rocking, followed by quenching with 0.125 M glycine for 5 min. Samples were centrifuged at 4000 rpm for 5 min at 4°C and washed three times in PBS supplemented with protease inhibitors and sodium butyrate to preserve histone acylations. Crosslinked tissue was homogenized in Covaris ChIP lysis buffer using a type B glass dounce (20 strokes per sample), followed by sequential washes and resuspension in SDS shearing buffer. Chromatin was sonicated for 10 cycles using a Diagenode Bioruptor Pico to generate DNA fragments of 200–500 bp. Following centrifugation at 10,000 × g for 5 min to remove insoluble debris, supernatants were collected, and 50% Protein G slurry was added for preclearing at 4°C for 30 min. For immunoprecipitation, chromatin was incubated overnight at 4°C with anti-pan Kcr antibody (PTM BIO, #PTM-501) or normal rabbit IgG (negative control; Diagenode, #C15410206) in ChIP dilution buffer. The next day, antibody–chromatin complexes were captured with blocked Dynabeads for 2 h at 4°C and subjected to sequential washes with low-salt, high-salt, LiCl, and TE buffers. Chromatin was eluted twice with 100 µL of ChIP elution buffer at 65°C and pooled. Crosslinks were reversed by 5 M NaCl and incubating at 65°C overnight, followed by RNAse A and proteinase K digestions. DNA was purified using the QIAquick PCR purification kit (Qiagen) and eluted in nuclease-free water.

### ChIP-Seq library preparation, sequencing, and analysis

ChIP-Seq libraries were prepared using KAPA library preparation kit. Indexed libraries were pooled and sequenced in illumina NovaSeq 6000. Paired-end sequencing was performed for all input and ChIP samples, generating approximately 100 million reads per sample with a read length of 100 nucleotides. ChIP-Seq reads were trimmed from both 5’ and 3’ ends using Trimmomatic paired end mode until the final base had a quality score > 30, discarding reads with < 23 bp. ChIP-Seq trimmed reads were aligned to mm10 reference genome using bwa aligner. Differential histone crotonylation between samples was assessed using the ‘MAnorm’ algorithm^77^ which quantitatively compares ChIP-seq signal intensities between two samples by normalizing against the set of peaks shared in both datasets. Broad peaks were first called with ‘MACS2’ using lenient thresholds to capture histone mark enrichment. These peak sets were then provided to MAnorm as input. The program merges peaks across both samples to define a unified reference, identifies overlapping peaks, and computes read densities for each region. MAnorm then models the relationship between samples by performing a robust linear regression on the log^2^ fold-change versus mean intensity of common peaks, thereby normalizing systematic differences in sequencing depth or background enrichment. For each peak, MAnorm reports a normalized M-value (log^2^ fold-change) and an associated p-value, which imply the degree and significance of differential enrichment. To focus on regulatory elements, significant differential peaks were filtered to retain only those located within ±2,000 bp of gene promoters (defined relative to the annotated TSS).

### Comparative analysis of learning-induced gene targets regulated by H3K27ac and Kcr

We compared gene targets of Kcr in dorsal hippocampus with gene promoters enriched with H3K27ac in the cortex. Raw sequencing dataset focusing on H3K27ac modifications in cortical neurons (GSM1629381, GSM1629397)^42^ were downloaded from the Sequence Read Archive using fastq-dump from sratoolkit. We then ran the script ‘ChIP-seq_SE_pipeline.sh’ for each raw dataset and normalized results by their respective ChIP-seq inputs using the script ‘subtractTwoWig.py’. Peak calling was performed using ‘MACS2’, and identified peaks were overlapped with gene annotations within a region spanning 2,000 base pairs upstream to 2,000 base pairs downstream of the transcription start site (TSS). We analyzed the resulting gene lists by overlapping them with learning-induced genes from our previously published RNA-seq study ^28^ to associate fold change values with different gene sets harboring distinct epigenetic marks.

### In situ hybridization (RNAscope)

*In situ* hybridization was performed using the RNAscope™ Multiplex Fluorescent Reagent Kit v2 (Advanced Cell Diagnostics) according to the manufacturer’s protocol as previously described^30^. Briefly, 20 μm cryosections obtained from fixed frozen brains were mounted on Superfrost™ Plus microscope slides (Fisherbrand). Slides then underwent serial dehydration in Ethanol, followed by Hydrogen Peroxide treatment, Target Retrieval, and Protease III treatment. Hybridization of probes was done at 40°C for 2 hr in an HybEZ oven using a probe against *Gria4*. The probe signal was amplified with Pre-amplifiers (AMP 1-FL, AMP 2-FL, and AMP 3-FL) and counterstained with OPAL 570 reagent (#FP1488001KT, Akoya Biosciences). Finally, coverslip mounting was done on the slides using ProLong™ Diamond Antifade Mountant with DAPI (ThermoFisher Scientific, P36962). The slides were stored in 4^0^C until they were imaged.

### Adeno-associated virus (AAV) constructs and stereotactic surgeries

AAV_9_-CaMKIIα-EGFP and AAV_9_-CaMKIIα-CDYL-V5 were purchased from VectorBuilder (VectorBuilder Inc). AAV_9_-CaMKIIα-GFP-Cre (#105551), pAAV_9_-hSyn-iGluSnFr (#98929), and pAAV_9_-hSyn1-GCaMP6s-P2A-mRuby3 (#112005) were purchased from Addgene. Mice were anesthetized using 5% isoflurane. A steady flow of 2.5% isoflurane was maintained throughout the remainder of the stereotactic surgery. 1 µl of respective AAVs were bilaterally injected into the dorsal hippocampus (coordinates: anteroposterior, −1.9 mm, mediolateral, ±1.5 mm, and 1.5 mm below bregma). Following viral infusion, drill holes were closed with bone wax (Lukens) and the incisions were sutured. Intraperitoneal (IP) injections of Meloxicam (5 mg/kg) were administered for 2 successive days after the surgery to manage the post-operative pain.

### Spatial object recognition (SOR) task

All the behavioral experiments were performed based on previously published studies^30,58^ during the light cycle in between Zeitgeber time (ZT) 0 through 2. Mice were individually housed for 7 days before the training. Animals were handled for 2 mins each day for 5 successive days before training. In the strong training paradigm, animals were habituated in an open field for 6 minutes. This was followed by three 6-minute sessions inside the same arena containing three different glass objects. These objects were placed at specific spatial coordinates with respective to a spatial cue within the arena. An inter-trial interval of 5 minutes was set in-between the three training sessions. In the sub-threshold training paradigm, mice were subjected to three 3-minute training trials in the open field with two objects and an inter-trial interval of 5 minutes in between sessions. 24 hr after training, the animals were returned to the arena with one of the objects displaced to a novel spatial coordinate. Exploration time around all the objects were then manually scored. Percent preference towards the displaced object was calculated using the following equation:

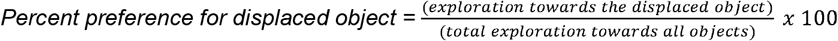

For the histone Kcr profiling and the single-nuclei experiments, animals were euthanized by cervical dislocation 1 hr after the last training trial in the SOR task. Hippocampal tissue was flash frozen and stored at -80^0^C. Home caged animals were euthanized within the same ZT window to eliminate the possible confounding effects of circadian rhythmicity. To examine the learning-induced expression of Kcr in hippocampal subregions, mice were perfused 1 hr after the training session, and whole brains were harvested for IHC.

### Contextual fear conditioning

Contextual fear conditioning was performed using a sub-threshold learning paradigm. In brief, mice were handled daily for five days before conditioning. The conditioning protocol involved a single 2-second, 0.75 mA footshock delivered 2.5 minutes after the mice were introduced into the chamber. Mice remained in the chamber for an additional 30 seconds before being returned to their home cage. 24-hrs later, they were reintroduced to the same chamber for 5 mins. Freezing behavior was assessed using FreezeScan software (CleverSys Inc.).

### Nuclei isolation

Nuclei isolation form frozen hippocampal tissue was performed according to the manufacturer’s protocol (Chromium Nuclei Isolation Kit with RNAse Inhibitor, 10x Genomics, #1000494). Briefly, frozen tissue was homogenized in 500 μl of Lysis Buffer using Dounce homogenizers. The lysate was transferred to a Nuclei Isolation Column and centrifuged at 16,000 x g for 30 seconds at 4^°^C. The pellet was then resuspended in 500 μl of Debris Removal Buffer and centrifuged at 700 x g for 10 minutes at 4^°^C. The nuclear pellet was then resuspended with 1 ml of Wash Buffer and centrifuged at 500 x g for 5 minutes at 4^°^C. Finally, the nuclear pellet was resuspended in 50 μl of Resuspension Buffer. Nuclei count was manually done using a Hemocytometer.

### Single-nuclei multiomic data processing and analysis

Raw sequencing data were processed using the ‘Cell Ranger ARC’ pipeline (v2.0.2) with the cell ranger-arc mm10 reference. Default parameters were used to align reads, count unique fragments or transcripts, and filter high-quality nuclei. HDF5 files for each sample (Saline1, Saline2, Crotonate1, Crotonate2) containing barcoded RNA counts and ATAC fragments per cell cluster were loaded into Seurat (Read10X_h5). This resulted in the generation of four Seurat objects, each containing both RNA and ATAC assays. Nuclei with outliers within the ATAC and RNA QC metrics (<200 and >100,000 ATAC read counts, <200 and >50,000 RNA read counts, nucleosomal signal□>□4, TSS enrichment□<□3, %reads in peaks < 15 and percentage of mitochondrial reads > 5) were removed.

To analyze the RNA component of the multiomics data, gene counts were normalized, and log transformed (LogNormalize). The top 2,000 most variable features that distinguish each cell were identified using ‘FindVariableFeatures’ (selection.method□=□’vst’). Features that are repeatedly variable across cells and datasets were selected for integration (‘SelectIntegrationFeatures’). We then identified anchors (‘FindIntegrationAnchors’’), which took the list of 4 individual Seurat objects for each sample as input. These anchors were used to integrate the four datasets together (‘IntegrateData’). Linear dimensionality reduction was performed on the integrated Seurat object by principal component analysis (runPCA, npcs□=□30). A k-nearest-neighbours graph was constructed based on Euclidean distance in PCA space and refined (‘FindNeighbors’), following which the nuclei were clustered using the Louvain algorithm (FindClusters, resolution□=□0.5). Clusters were visualized with UMAP (runUMAP, dims□=□30). Both RNA and ATAC assays were used to identify cell-type specific signatures of biomarkers. Differentially expressed genes (DEGs) in individual clusters between saline and crotonate treated groups were calculated (FindMarkers, test.use□=□’wilcox,’ Padj□<□0.05, absolute logFC.threshold of 0.2).

To analyze the ATAC component of the multiomics data, the default assay was switched to ATAC prior to integrating the four Seurat objects, and peak calling was performed. The set of peaks identified by ‘Cellranger’ often merges distinct peaks that are close together - confounding the motif enrichment analysis and peak-to-gene linkage. We were able to circumvent this concern and identify a more accurate set of peaks by using the ‘MACS2’ (CallPeaks) peak calling feature on all cells together. Peaks on nonstandard chromosomes and in genomic blacklist regions were removed (‘keepStandardChromosomes’ and ‘subsetByOverlaps’). A frequency-inverse document frequency normalization was performed across cells and peaks (‘RunTFIDF’). Thereafter, a feature selection was performed using all the peaks as input (FindTopFeatures, min.cutoff = 5). The selected peaks went through a dimensional reduction on the TF-IDF normalized matrix using a singular value decomposition (‘RunSVD’). To identify the differentially accessible regions (DAR) between crotonate versus saline group, the Seurat function ‘FindMarkers’ was used using logistic regression (LR) as a method to test significance. The DARs with adjusted p value < 0.05 and absolute log2foldchange threshold of above 0.2 were considered as significantly differentially accessible. Further, the DARs were annotated using ‘Closestfeature’ function from Signac package as well as ‘annotatePeak’ function from ‘ChIPseeker’ package. The DEGs were furthered correlated with DARs where the DARs were selected only from the promoter (+/-2kb from transcription start site) and genebody regions, filtering out the peaks from distal genomic regions and downstream of 3’UTR^78^. The genes found concordant in both DEG and DAR lists were shortlisted for downstream gene ontology enrichment analysis. UpSet plots were generated using UpSetR package.

### Gene Ontology enrichment analysis

The concordant DARs and DEGs were analyzed for Molecular Function (MF) enrichment analysis by using the ‘clusterProfiler’ package with the default criteria (pvalueCutoff = 0.01 and qvalueCutoff = 0.05). Here, the significant upregulated DEGs (adj p value < 0.05 and log2FC > 0.2) concordant with the significant more accessible DARs (adj p value < 0.05 and log2FC > 0.2) were used to generate the upregulated gene list, whereas the significant downregulated DEGs (adj p value < 0.05 and log2FC < -0.2) concordant with the significant less accessible DARs (adj p value < 0.05 and log2FC < -0.2) were used to generate the downregulated gene list. The DARs were selected only from the promoter (+/-2kb from transcription start site) and gene body regions, filtering out the peaks from distal genomic regions and regions downstream of 3’UTR. All further data visualizations were made using clusterProfiler package.

### TF motif enrichment analysis

To identify cell-type specific regulatory sequences, we performed transcription factor motif enrichment analysis on the DEGs that were found to have concordant differentially accessible peaks in the promoter region. Here, we restricted TF motif enrichment only in the promoter region (a window of 2000bp upstream and downstream of transcription start site). Motif enrichment was performed using ‘FindMotifs’ function of signac package. The motifs that had adjusted p value < 0.05 were considered significant. The top 15 significant motifs from the clusters were plotted as heatmap using ComplexHeatmap package. Weight matrices for the top motifs were also plotted to visualize the motif sequences.

### Cell-cell communication analysis

The cell-cell communication analysis on the snRNA-seq data was performed using CellChat (v2.1.2). The Saline and Crotonate RNA normalized counts were taken and individual cellchat objects were created. Ligand-receptor (LR) interactions in each group were calculated by identifying overexpressed ligands or receptors with a log fold change cutoff 0.1 (identifyOverExpressedGenes(thresh.fc = 0.1, thresh.p = 0.05)) followed by identifying interactions if LR pairs are overexpressed (identifyOverExpressedInteractions). To assign each interaction with a probability score, computeCommunProb function was used with a default statistical method called ‘trimean’. Cell-cell communication was filtered to have minimum cell number 10 in each cell group (filterCommunication(min.cells = 10)). After that communication probability on signaling pathway level was calculated by summarizing the communication probabilities of all LR interactions associated with each signaling pathway. Finally, the cell-cell communication network was aggregated by counting the number of links or summarizing the communication probability amongst the cell groups. Thereafter, the cellchat objects were merged and compared using netVisual_aggregate, netAnalysis_signalingChanges_scatter, and netVisual_chord_gene functions.

### Acute brain slice preparation

Acute hippocampal slices were prepared as described in published studies^52,79^. 2-4-month-old male mice were deeply anaesthetized with isoflurane, following which their brains were harvested for slice preparation. Coronal sections were obtained in ice-cold dissection buffer (in mM: 220 sucrose, 3.3 KCl, 5 MgSO_4_, 1.25 NaH_2_PO4, 25 NaHCO_3_, 11 D-glucose, and 0.2 CaCl_2_) and sections containing dorsal hippocampal representations were allowed to recover in a chamber containing normal artificial CSF (aCSF) (in mM; 116 NaCl, 3.3 KCl, 1.3 MgSO_4_, 1.25 NaH_2_PO4, 25 NaHCO_3_, 11 D-glucose, 1.3 CaCl_2_) saturated with carbogen (95%O_2_ / 5%CO_2_) at 32°C. The slices were incubated for at least 2 hrs either in normal aCSF or 5 mM Crotonate before being transferred to a submerged chamber for imaging, constantly supplied with aCSF (flow rate: 2.5 ml/min) also maintained at 32°C. For the experiment described in **Fig. S9**, the slices were incubated for 5-10 mins with the drug, following which they were transferred to the submerged chamber containing aCSF for imaging.

### Mesoscopic imaging and image analysis

Acute coronal sections containing hippocampi transduced with either iGluSnFr or GCaMP6s were placed in a submerged chamber attached to the stage of an upright microscope (Olympus MVX10), continuously perfused with aCSF saturated with carbogen and maintained at 32-34°C. A monopolar stainless steel stimulation electrode (AM systems) was placed on in the Stratum Radiatum to stimulate the Schaffer Collateral pathway. Optical excitation was delivered by an LED light (X-Cite XYLIS XT720S). Fluorescent mesoscopic imaging of the entire brain slice was performed using a 2x objective (NA 0.5). The LED light was band-passed through a 470/40 nm filter and a dichroic 495 nm filter, while the emitted light was isolated using a 525/50 nm filter. For stimulus trains, 10 stimuli (3mV, 500 µs) were delivered with variable inter-stimulus intervals (for 10 Hz: 100ms; for 20Hz: 50ms; for 50Hz: 20ms), as done in previous studies^52^. Images were acquired with a CMOS camera (Basler Ace acA4112-30µm) at a frame rate of 10 Hz (GCaMP6s imaging) or 90-100 Hz (iGluSnFr imaging) with 4:1 binning. The camera acquisition and LED excitation were digitally synchronized using Micro-manager (ImageJ). Raw images in time series were background-subtracted and smoothed (median filter). Time series were baseline subtracted, scaled to baseline means (ΔF/F_0_, baseline: ∼2s before the stimulation / response onset), and then compressed into Max Intensity Projections (MIPs). For Glutamate transients (iGluSnFr), responsive regions (ROIs) were generated by converting the ΔF/F_0_ MIPs into binary (Threshold: mean + SD). For calcium transients, hippocampal area that showed ectopic expression of GCaMP6s was selected as the ROI. Fluorescence traces generated using the respective ROIs were analyzed in MATLAB (MathWorks). Ca^2+^/ Glutamate response parameters (peak amplitude, locations, width, prominence) were determined using the ‘FindPeaks’ function in MATLAB (minimum peak prominence: 0.025 for Glutamate, 0.05 for Ca^2+^). Area under the curve (AUC) was then calculated using trapezoidal numerical integration (‘Trapz’ function) in MATLAB. For glutamate transients, the slow and fast components of the decay kinetics (Tau) were determined by fitting two single-term exponential fits to the decay of the response (τ_fast_: stimulus offset to ∼4s, τ_slow_: ∼4s to 10s).

### Statistics

Behavioral and biochemical data were analyzed using unpaired two-tailed t-tests, one-way ANOVA, or two-way ANOVAs (sometimes with repeated measures as the within-subject variable). Sidak’s multiple comparison tests or Dunnett’s multiple comparison tests were used for post-hoc analyses wherever required. In the iGluSnFR and GCaMP6s imaging data, normality of the distributions was assessed using the D’Agostino–Pearson K2, Anderson-Darling tests, and QQ plots. If data was log-normal, a Lognormal Welch’s t test was used. Outliers were detected using Grub’s or IQR tests (for parametric and nonparametric data), although no outliers were detected or excluded. In all the experiments, differences were considered statistically significant when p<0.05. As indicated for each figure panel, all data were plotted in box plots.

### Ethics

All procedures on mice in this study were conducted according to US National Institutes of Health guidelines for animal care and use and were approved by the Institutional Animal Care and Use Committee of the University of Iowa, Iowa.

### Reporting summary

Further information on research design is available in the Nature Portfolio Reporting Summary linked to this article.

## Acknowledgments

We thank the Neural Circuits and Behavior Core and the Iowa Institute of Human Genetics (IIHG) core for using their facilities, Emily N. Walsh, Lucy Langmack, Quinlan Truax, Adam J Rauckhorst, and Rebekah M. Peplinski for their technical assistance, and K. Peter Giese and Jacob Michaelson for their valuable input on this work.

## Author contributions

S.C. and T.A. conceived the study. S.C. and U.M. designed the experiments, interpreted the results, and wrote the article with inputs from T.A. and E.B.T. U.M., S.C., and S.E.B. performed the behavioral tasks, stereotactic surgeries, drug treatment, and biochemical experiments. S.G. and N.R. assisted with the analysis of behavioral data and biochemical experiments. E.B.T. provided the ACSS2 mice. B.B. performed the bioinformatic analyses with inputs from Y.V. P.S performed the functional epifluorescence experiments with inputs from J.G. All the authors edited the manuscript.

## Data availability

The datasets generated in this study have been deposited in the NCBI Gene Expression Omnibus (GEO) database under accession code GSE281007. Sequencing files for the single-nuclei multiomics (RNA-seq + ATAC-seq) and ChIP-Seq have been made publicly available through GSE281007.

## Code availability

The code used for the analyses to generate the figures related to Single-nuclei multiomics data can be accessed through GitHub (https://github.com/ChatterjeeEpigenetics/CrotonylationMultiomics2024).

