## Supplemental information for "Lysine Crotonylation Acts as an Epigenetic Switch for Glutamate Neurotransmission and Spatial Memory"

**Running title:** Histone crotonylation regulates memory storage and glutamate signaling.

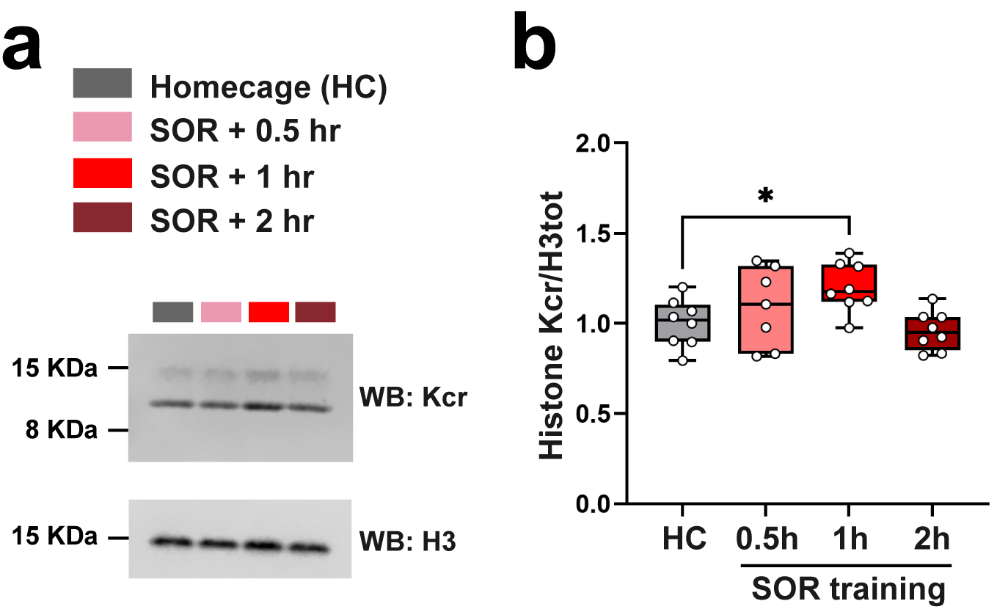

**Supplementary Figure 1. Learning-dependent increase in histone lysine crotonylation in the dorsal hippocampus. a.** Western blot showing histone crotonylation analyzed from the dorsal hippocampus of mice trained in spatial object recognition (SOR) task and euthanized at 0.5, 1, or 2h after training. Homecage (HC) mice were used as controls. **b.** Quantification of **a.** One-way ANOVA: Kcr: F(3, 27)=3.953, p= 0.0185. Dunnett multiple comparisons tests: *p=0.0379 (HC versus 1 hr). Homecage (n = 8), SOR + 0.5 hr (n = 7), SOR + 1 hr (n=8), SOR + 2 hr (n=8), males only. All box plots: the center line represents the median, the box edges represent the top and bottom quartiles (25th to 75th percentiles), and the minimum and maximum whiskers.

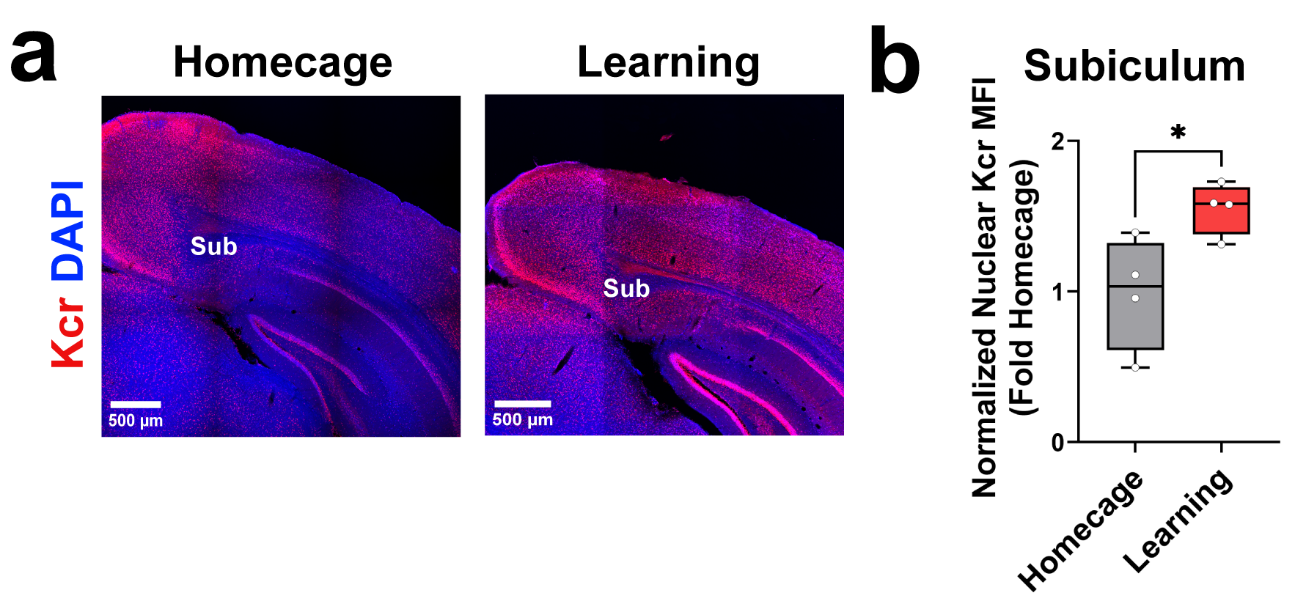

**Supplementary Figure 2. Spatial learning elevates lysine crotonylation levels in the Subiculum. a.** Immunofluorescence using anti-Kcr antibody showing levels of Kcr in Subiculum (Sub) of HC and learning (SOR+1hr) mice. **b.** Quantification of **a.** Unpaired t test: t (6) = 2.733, *p = 0.034. Homecage (n = 4) and SOR+1 hr (n = 4), males only. All box plots: the center line represents the median, the box edges represent the top and bottom quartiles (25th to 75th percentiles), and the minimum and maximum whiskers.

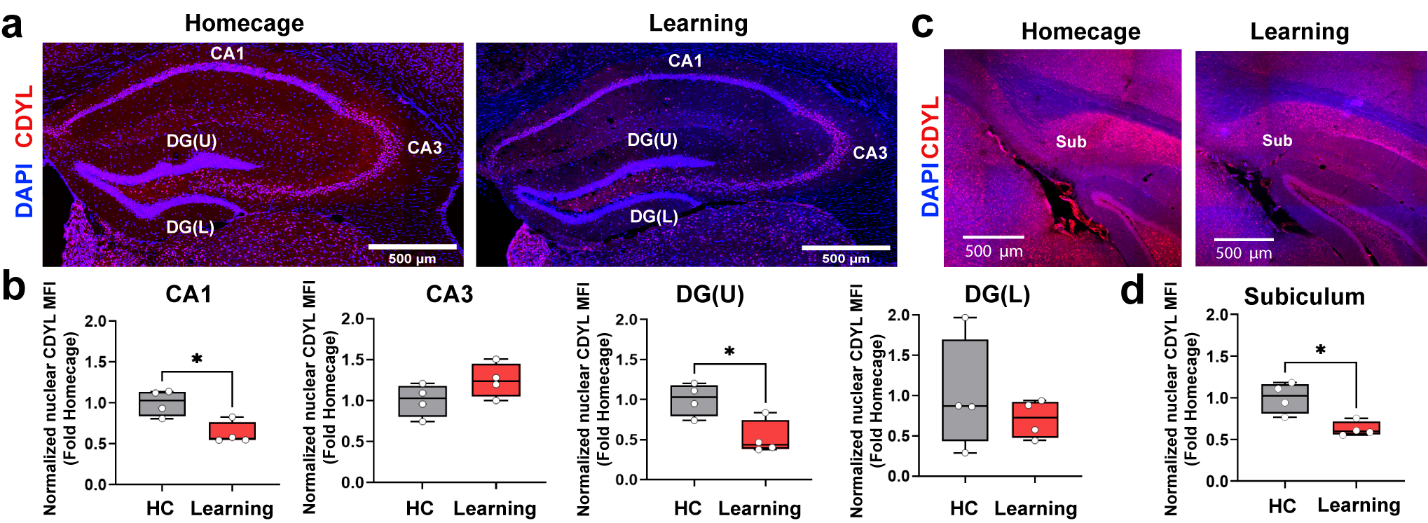

**Supplementary Figure 3. Spatial learning reduces hippocampal CDYL levels. a-d.** Immunofluorescence using an anti-CDYL antibody was performed on brain slices from mice 1 hr after training in the SOR task. Hippocampal sub-regions CA1 (**3a**), CA3 (**3a**), DG upper blade (U) (**3a**), DG lower blade (L) (**3a**), and Subiculum (**3c**) were examined. Normalized Mean Fluorescent Intensity (MFI) of nuclear CDYL levels across the groups. **b, d.** Quantification of **a** and **c** respectively. For CA1 (**3b**): Unpaired t test: t (6) = 3.598, *p = 0.0114. Homecage (n = 4) and SOR+1 hr (n = 4), males only. For CA3 (**3b**): Unpaired t test: t (6) = 1.704, p = 0.1393. Homecage (n = 4) and SOR+1 hr (n = 4), males only. For DG upper blade (**3b**): Unpaired t test: t (6) = 3.254, *p = 0.0174. Homecage (n = 4) and SOR+1 hr (n = 4), males only. For DG lower blade (**3b**): Unpaired t test: t (6) = 0.7837, p = 0.463. Homecage (n = 4) and SOR+1 hr (n = 4), males only. For Subiculum (**3d**): Unpaired t test: t (6) = 3.657, *p = 0.0106. Homecage (n = 4) and SOR+1 hr (n = 4), males only. All box plots: the center line represents the median, the box edges represent the top and bottom quartiles (25th to 75th percentiles), and the minimum and maximum whiskers.

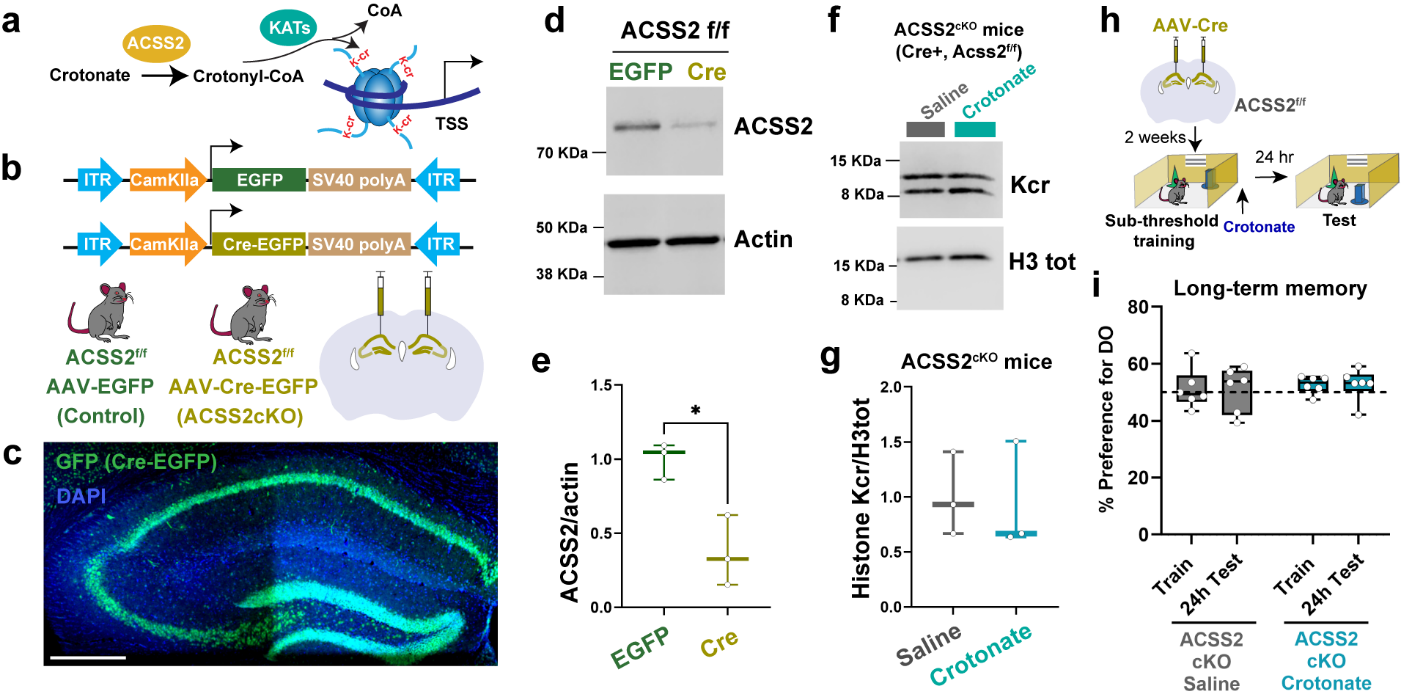
**Supplementary Figure 4. Crotonate-mediated memory enhancement is dependent on ACSS2*.* a.** Schematic of ACSS2 in regulating histone crotonylation. **b.** Schematic of viral constructs used to conditionally knock down ACSS2 in excitatory neurons of dorsal hippocampus. AAV_9_-CaMKIIα-EGFP served as vector control and AAV_9_-CaMKIIα-Cre-EGFP was used to drive expression of Cre recombinase in excitatory neurons of ACSS2^f/f^ mice. **c.** Immunofluorescence image using GFP antibody showed expression of the Cre-EGFP in the dorsal hippocampus two weeks following viral infusion. **d-e.** Western blot from whole cell extracts of the dorsal hippocampus of ACSS2^f/f^ mice infused with AAV-EGFP or AAV-Cre-EGFP. Normalized band intensity of ACSS2 across the groups: Unpaired t test: t (4) = 4.118, *p = 0.0146. EGFP (n = 3) and ACSS2 (n = 3), males only. **f-g.** Western blot using Kcr antibody from core histones extracted from the dorsal hippocampus of ACSS2 cKO mice (ACSS2^f/f^ infused with AAV-Cre-EGFP) administered either crotonate (200 mg/kg, oral gavage) or saline immediately after SOR training. Normalized band intensity of pan Kcr across the groups: Unpaired t test: t (4) = 0.1853, p = 0.862. Saline (n=3), Crotonate (n=3), males only. **h-i.** Crotonate treatment after training using a sub-threshold SOR learning paradigm does not enhance long-term memory in ACSS2 cKO mice. Two-way ANOVA: no significant session x treatment interaction F_(1, 10)_ = 0.0007554, p= 0.9786. Saline (n=6), Crotonate (n=6), males only. All box plots: the center line represents the median, the box edges represent the top and bottom quartiles (25th to 75th percentiles), and the minimum and maximum whiskers

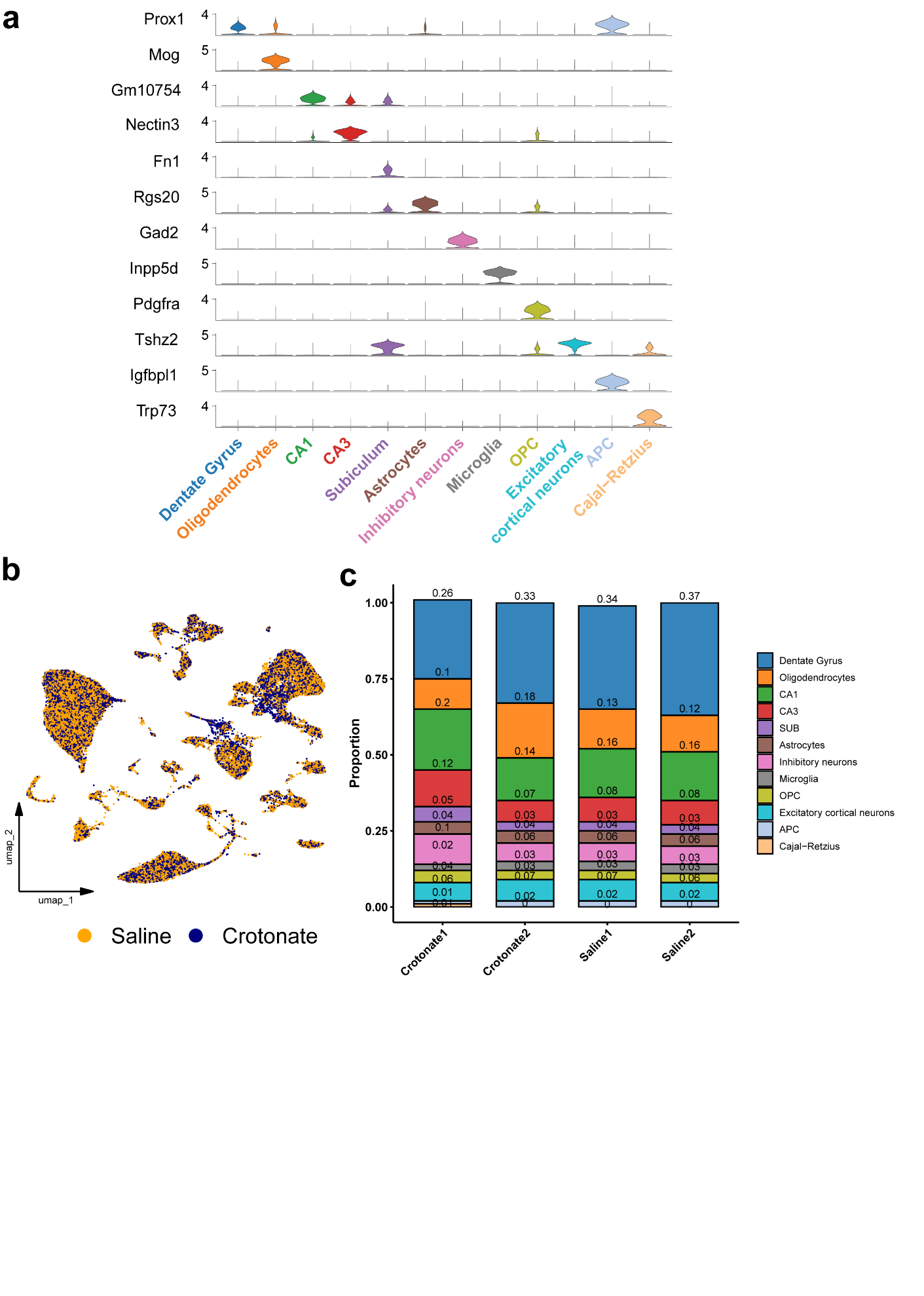

**Supplementary Figure 5. a.** Violin plot showing the expression profiles of marker genes across different cell types of the dorsal hippocampus. **b.** UMAP plots depicting cell-type specific clusters in saline-treated and crotonate-treated groups. **c.** Cell-type-proportions for each sample across experimental groups.

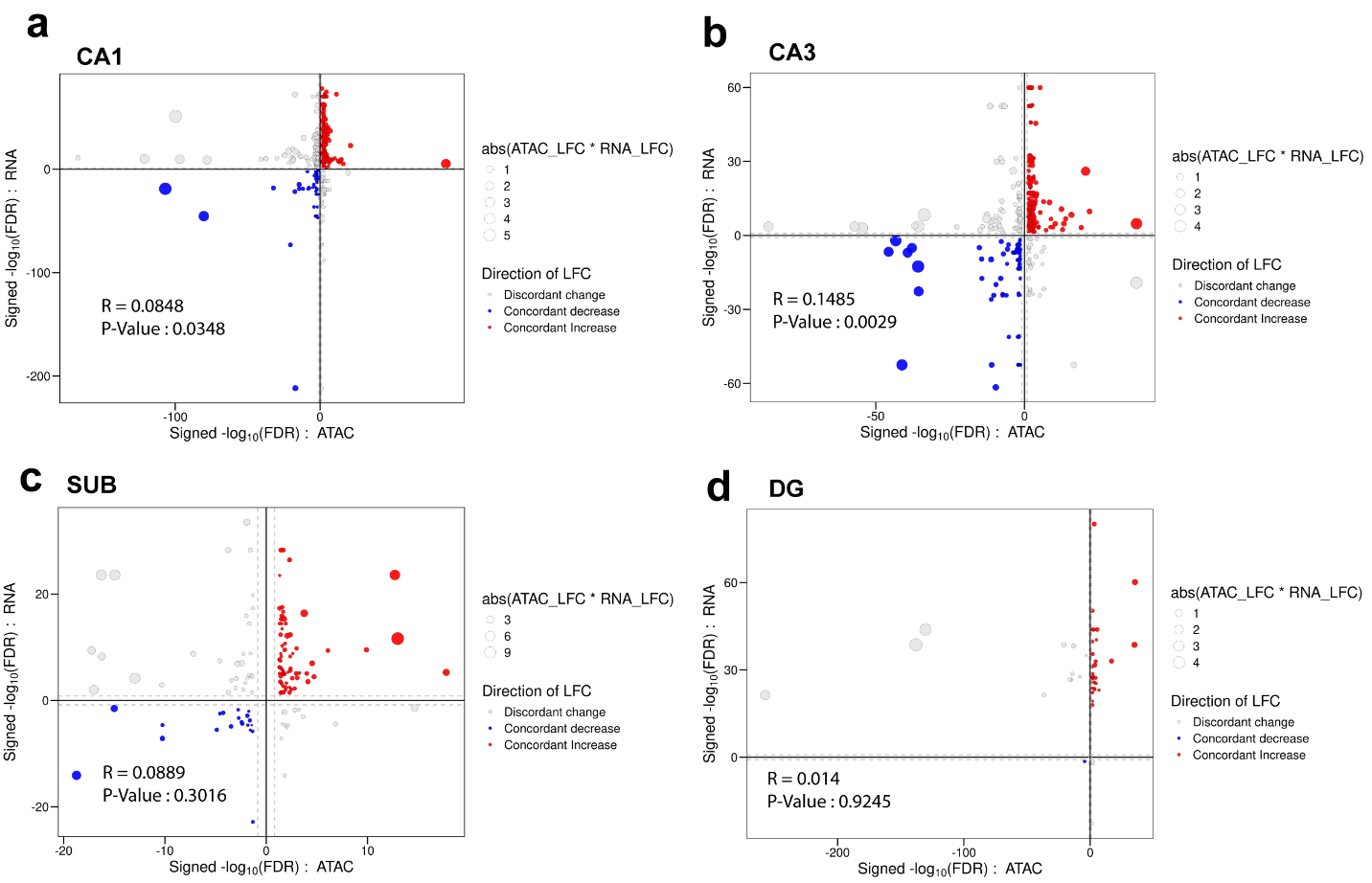
**Supplementary Figure 6. Quadrant plots depicting the correlation between genes with significant differential expression and genes with significant DARs in hippocampal principal neuronal layers. a.** CA1, **b.** CA3, **c.** Subiculum. **d.** Dentate gyrus (DG).

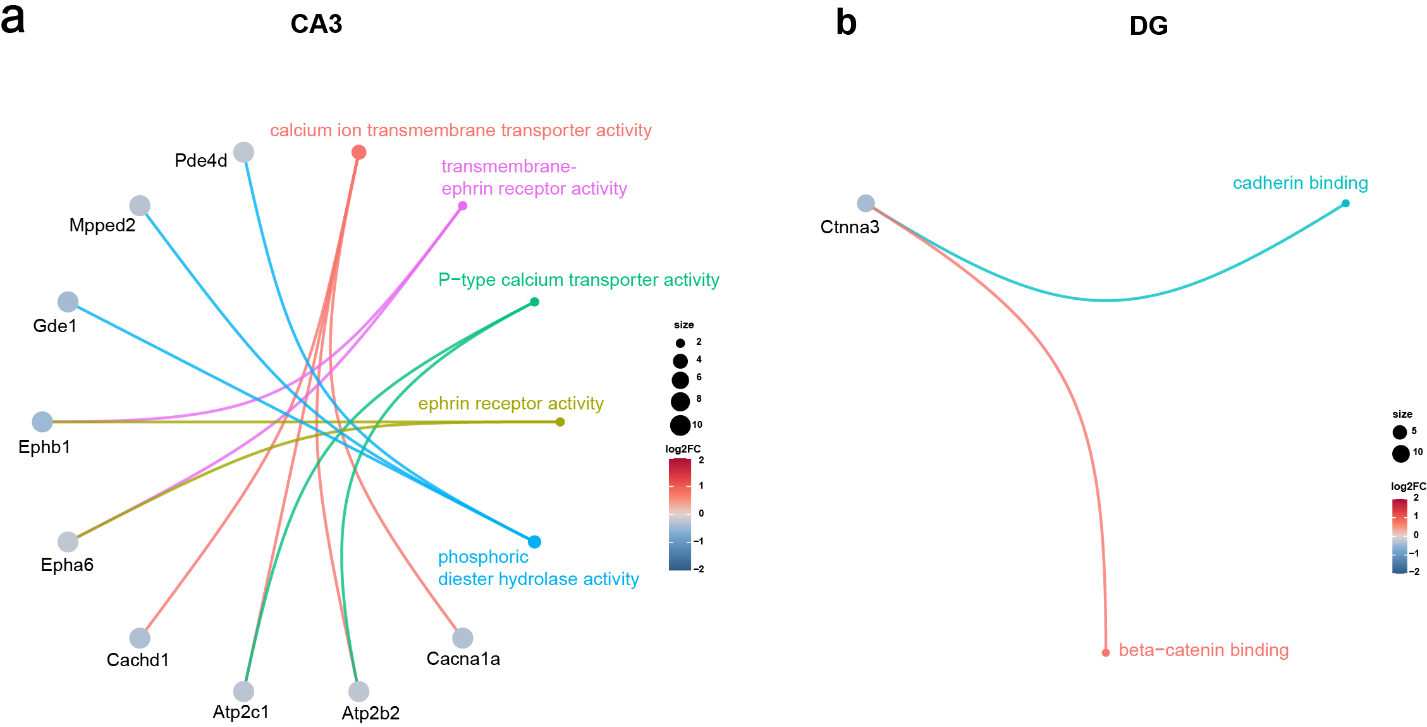

**Supplementary Figure 7. Top 5 most significantly enriched GO: Molecular function patterns from downregulated genes. a.** CA3. **b.** Dentate gyrus (DG).

**Supplementary**
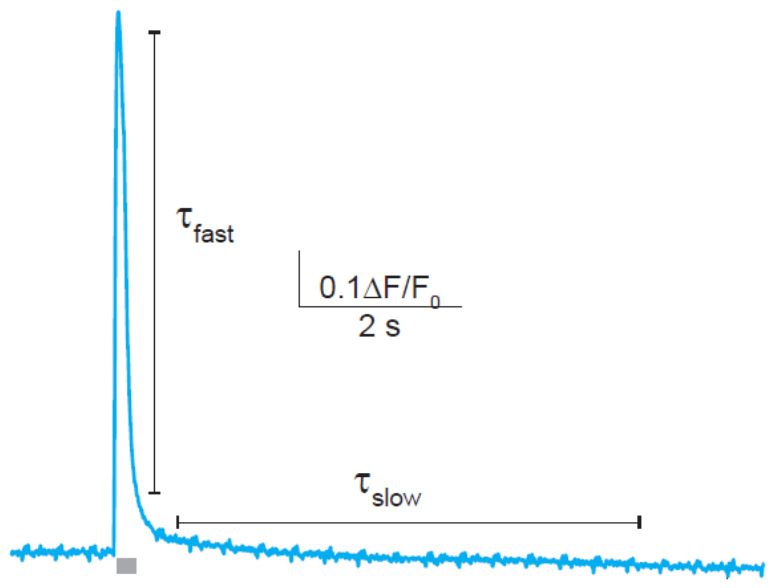
**Figure 8. Computation of glutamate clearance rates.** Glutamate clearance kinetics were determined by fitting the decay of the glutamate response with two single-term exponential equations (black solid line); blue, glutamate fluorescence from a single trial; τ_fast_: stimulus offset to ~4s, τ_slow_: ~4s to 10s.

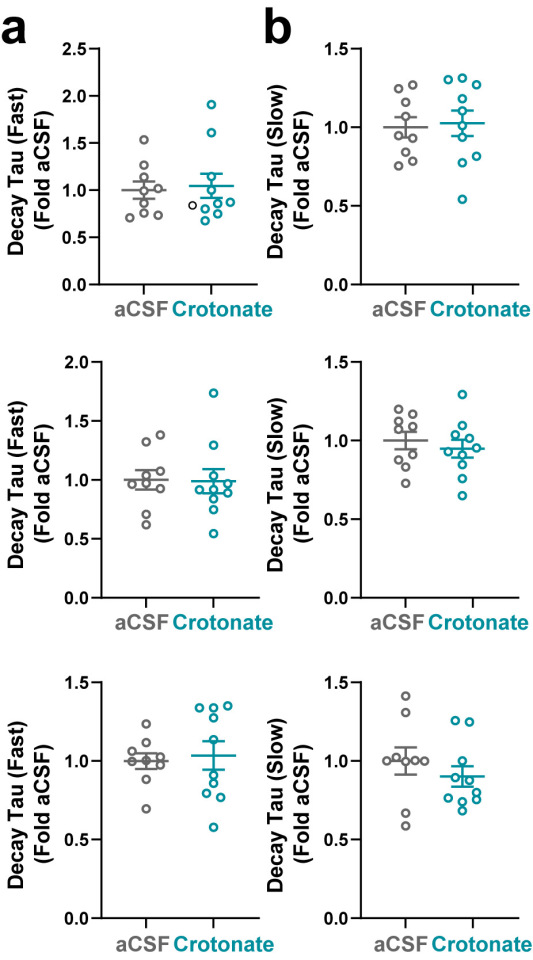

**Supplementary Figure 9. Crotonate-dependent increases in Kcr does not affect glutamate decay kinetics. a.** No differences in fast decay kinetics were observed between the two groups across all stimulation frequencies. For 10 Hz (top): Lognormal Welch’s t test: t (16.74) = 0.1425, p = 0.884. aCSF (n = 9) and Crotonate (n = 10). For 20 Hz (middle): Lognormal Welch’s t test: t (16.94) = 0.2031, p = 0.8415. aCSF (n = 9) and Crotonate (n = 10). For 50 Hz (bottom): Welch’s t test: t (13.89) = 0.3411, p = 0.7381. aCSF (n = 9) and Crotonate (n = 10). **b**. No differences in slow decay kinetics were observed between the two groups across all stimulation frequencies. For 10 Hz (top): Lognormal Welch’s t test: t (15.98) = 0.0768, p = 0.9397. aCSF (n = 9) and Crotonate (n = 10). For 20 Hz (middle): Welch’s t test: t (16.98) = 0.6512, p = 0.5236. aCSF (n = 9) and Crotonate (n = 10). For 50 Hz (bottom): Lognormal Welch’s t test: t (14.94) = 0.7948, p = 0.4392. aCSF (n = 9) and Crotonate (n = 10). All scatter plots: the center line represents the mean, error bars represent ±SEM.

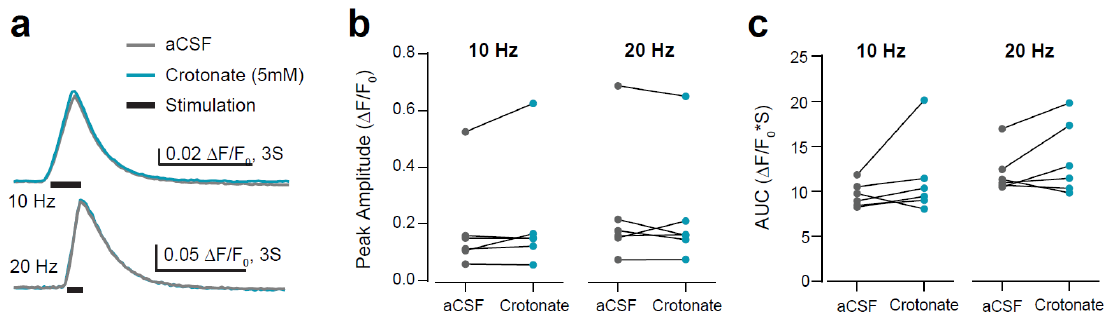

**Supplementary Figure 10. Examining non-genomic effects of crotonate.** Brief incubation with crotonate (5mM) does not exert any effect on the peak amplitude (**b**) or AUC (**c**) of stimulus-evoked calcium transients. **b.** Peak amplitude: For 10 Hz (left): Paired t test: t (5) = 1.446, p = 0.2077. For 20 Hz (middle): Paired t test: t (5) = 0.6143, p = 0.5659. **c.** Area under the curve (AUC): For 10 Hz (left): Paired t test: t (5) = 1.290, p = 0.2534. For 20 Hz (middle): Paired t test: t (5) = 1.522, p = 0.1886. aCSF (n = 6) and Crotonate (n = 6).
